# Nightmare or delight: taxonomic circumscription meets reticulate evolution in the phylogenomic era

**DOI:** 10.1101/2023.03.28.534649

**Authors:** Ze-Tao Jin, Richard G.J. Hodel, Dai-Kun Ma, Hui Wang, Guang-Ning Liu, Chen Ren, Bin-Jie Ge, Qiang Fan, Shui-Hu Jin, Chao Xu, Jun Wu, Bin-Bin Liu

## Abstract

Phylogenetic studies in the phylogenomics era have demonstrated that reticulate evolution greatly impedes the accuracy of phylogenetic inference, and consequently can obscure taxonomic treatments. However, the systematics community lacks a broadly applicable strategy for taxonomic delimitation in groups identified to have pervasive reticulate evolution. The red-fruit genus, *Stranvaesia*, provides an ideal model for testing the effect of reticulation on generic circumscription when hybridization and allopolyploidy define a group’s evolutionary history. Here, we conducted phylogenomic analyses integrating data from hundreds of single-copy nuclear (SCN) genes and plastomes, and interrogated nuclear paralogs to clarify the inter/intra-generic relationship of *Stranvaesia* and its allies in the framework of Maleae. Analyses of phylogenomic discord and phylogenetic networks showed that allopolyploidization and introgression promoted the origin and diversification of the *Stranvaesia* clade, a conclusion further bolstered by cytonuclear and gene tree discordance. The well-inferred phylogenetic backbone revealed an updated generic delimitation of *Stranvaesia* and a new genus, *Weniomeles*, characterized by purple-black fruits, trunk and/or branches with thorns, and fruit core with multilocular separated by a layer of sclereids and a cluster of sclereids at the top of the locules. Here, we highlight a broadly-applicable workflow for inferring how analyses of reticulate evolution in phylogenomic data can directly shape taxonomic revisions.

## 1. Introduction

Evolutionary biologists have aimed to reconstruct phylogenetic relationships between organisms using multiple lines of evidence, such as phenotypic and molecular characters (Mindell, 2013). Historically, bifurcating phylogenetic trees have been the most tractable model for representing the evolutionary histories of large taxonomic groups (Huson and Bryant, 2006; Mindell, 2013), with monophyly widely accepted as the gold standard for defining groups of taxa. Phylogenetic and phylogenomic data have been widely used to infer phylogeny among species, make taxonomic revisions, and understand the origin of life. Although phylogenetic trees are suitable for reflecting speciation, the strict bifurcating structure of trees limits their use in describing more complex evolutionary scenarios, such as hybridization, incomplete lineage sorting (ILS), and/or allopolyploidy (Huson and Scornavacca, 2011; Morrison, 2014). Increasing evidence has shown that reticulate processes such as hybridization and polyploidization promoted the diversification of many lineages, particularly angiosperms (Mallet, 2005; Rothfels, 2021), e.g., nearly one-third of extant vascular plants are estimated to be of polyploid origin (Wood et al., 2009). The genomic age has made it clear that reticulate evolution is pervasive in the tree of life, and a strictly bifurcating phylogeny is rarely the best representation of evolution (Morales-Briones et al., 2021; Cooper et al., 2022; Debray et al., 2022; Smith et al., 2022; Zhao et al., 2022). Many different taxonomic hierarchies have been rearranged in the past decades as we have gained access to more genomic data and inference methods, such as for the flowering plants (APG: Byng et al., 2016), pteridophytes (PPG: Schuettpelz et al., 2016), birds (Jarvis et al., 2014), and mammals (Upham et al., 2019).

Phylogenetic networks are an excellent method for representing reticulate evolutionary processes (Debray et al., 2022); this method differs from bifurcating phylogenetic trees by modeling numerous linked networks, and adding hybrid nodes (nodes with two parents) instead of allowing only nodes with a single ancestor (Arenas et al., 2008). The development of related tools for evolutionary network reconstruction has also greatly facilitated their wide use in recent studies of evolution (Huson and Bryant, 2006; Schliep et al., 2017; Solís-Lemus et al., 2017). However, the downstream impact of modeling hybridization and other reticulate processes on taxonomic treatments is understudied, and further investigation of the impact of reticulate evolution on taxonomy is needed. Additional study of angiosperms, and other clades with histories of frequent reticulation, are needed to characterize the strengths and limitations of these approaches when using phylogenomic datasets. Furthermore, the implications of how diagnosing and quantifying reticulate evolution impact systematic and taxonomic revisions need further study in the phylogenomic age.

One clade characterized by both reticulate evolution and taxonomic uncertainty, *Stranvaesia* Lindl., is placed in the Rosaceae, a plant family with extensive whole genome duplication (WGD, Xiang et al., 2017; Morales-Briones et al., 2021) and hybridization events (Liu et al., 2019, 2020a, 2020b, 2022; Hodel et al., 2021; Su et al., 2021). Robertson et al. (1991) summarized the intergeneric hybrids in the apple tribe Maleae (formerly as subfamily Maloideae; his Fig. 1), and nearly 15 out of 24 genera have been involved in hybridization. A recent phylogenomic study (Hodel et al., 2021) successfully explained the observed conflict in the peach subfamily, and elucidated the hybrid origin of the Maleae-Gillenieae clade (i.e., the wide hybridization hypothesis between distantly related tribes in the subfamily Amygdaloideae) using phylogenetic network analyses. Moreover, their study also detected the pervasive nuclear gene tree-species tree conflict and/or cytonuclear conflict in the subfamily Amygdaloideae of Rosaceae. These frequent reticulation events in Maleae obscure diagnostic characters among genera and present significant challenges to generic circumscription. Historically, *Stranvaesia* has been either treated as a separate genus (Roemer, 1847; Decaisne, 1874; Wenzig, 1883; Focke, 1888; Koehne, 1890, 1891; Rehder, 1940, 1949; Yu and Ku, 1974; Lu and Spongberg, 2003) or merged into *Photinia* Lindl. (Lu et al., 1990, 1991; Li et al., 1992; Zhang, 1992). The red-fruit genus *Stranvaesia* and its allies thus represent a good case study for exploring the taxonomic treatment of a lineage with frequent hybridization and/or allo/autopolyploidy.

**Fig. 1.**
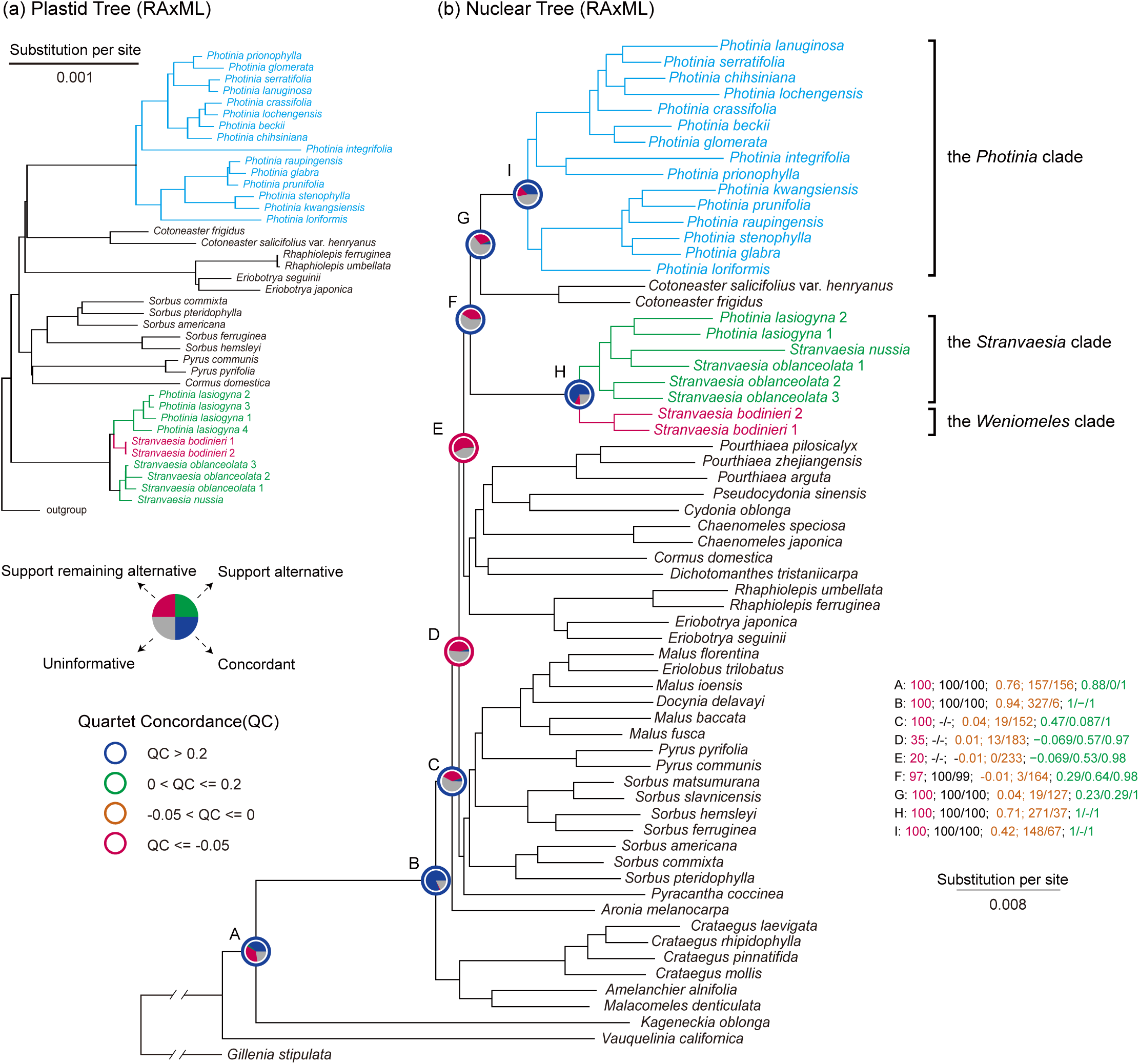
(a) A portion of the RAxML tree of *Stranvaesia* in the context of Maleae using the 78 concatenated plastid coding sequences (CDSs) supermatrix, see Fig. S2 for the whole tree. (b) Maximum Likelihood (ML) tree of *Stranvaesia* within Maleae inferred from RAxML analysis using the concatenated 426 single-copy nuclear genes (SCNs) supermatrix. Pie charts on the nine nodes (A-I) represent the proportion of gene trees that support that clade (blue), the proportion that support the main alternative bipartition (green), the proportion that support the remaining alternatives (red), and the proportion (conflict or support) that have < 50% Bootstrap Support (BS, gray). All the other pie charts in detail refer to Fig. S10. The color of the circle around the pie chart represents the value range of Quartet Concordance (QC), where QC > 0.2 is painted in dark blue, 0 < QC ≤ 0.2 is painted in light green, −0.05 < QC ≤ 0 is painted in organge, and QC ≤ −0.05 is painted in red. The QC value of the other nodes refers to Fig. S11. The numbers (bottom right) indicate values associated with those nodes; they are BS values estimated from RAxML analysis (e.g., A: 100 labeled by red; see Fig. S7 for all nodes BS), the SH-aLRT support and Ultrafast Bootstrap (UFBoot) support estimated from IQ-TREE2 (e.g., A: 100/100 labled by black; see Fig. S8 for all nodes support), the Internode Certainty All (ICA) score, the number of gene trees concordant/conflicting with that node in the nuclear topology estimated from *phyparts* (e.g., 0.76; 157/156 labled by organge; see Fig. S10 for all nodes), and Quartet Concordance/Quartet Differential/Quartet Informativeness estimated from Quartet Samping analysis (e.g., 0.88/0/1 labled by green; see Fig. S11 for all scores).

Traditionally, phylogenetic analyses use fragments of certain plastid and/or nuclear genes to reconstruct phylogenies, but these trees often reflect the history of a subset of genes rather than the history of the species (Zou and Ge, 2008). Phylogenomic approaches pave a promising path for utilizing genomic data to clarify relationships between species, infer rapid radiations and hybridization events in diverse lineages, and reconstruct the evolutionary history of organisms (Wen et al., 2013, 2017; Wickett et al., 2014). For example, Deep Genome Skimming (DGS, Liu et al., 2021) has often been used to target the high-copy fractions of genomes (Straub et al., 2012; Zimmer and Wen, 2015), including plastomes, mitochondrial genomes (mitogenomes), and nuclear ribosomal DNA (nrDNA) repeats, as well as single-copy nuclear genes (SCNs). Genome-level datasets often result in phylogenies, which even when strongly supported, have extensive underlying conflict, suggesting that in many cases, a network approach may better reflect true evolutionary relationships (Solís-Lemus et al., 2017; Wen et al., 2018). Liu et al. (2022) proposed a general procedure for inferring phylogeny and untangling the causes of gene tree and cytonuclear conflict; this pipeline can integrate multiple sources of sequencing data previously sequenced for one study, e.g., transcriptomic (RNA-Seq), genome resequencing (WGS), target enrichment (Hyb-Seq), and DGS data. Recently, more pipelines have also been proposed for modeling reticulate evolution and explicitly incorporating it into phylogenomic studies (Rose et al., 2020; Debray et al., 2022). However, in the phylogenomic era, taxonomic circumscription and the subsequent consequences on nomenclature have not been thoroughly investigated in lineages with frequent reticulation, such as hybridization and allopolyploidization events. This study aims to develop a pipeline for making robust taxonomic delimitations with guidance from phylogenetic network results. Here, using the red-fruit genus *Stranvaesia* as a test case, we propose a workflow that explicitly considers reticulate evolution in the taxon delimitation process. This workflow is demonstrated using data from *Stranvaesia* and its relatives to show its broad applicability across the Tree of Life (ToL).

## 2. Materials and Methods

### 2.1. Taxon sampling, DNA extraction, and sequencing

To clarify the phylogenetic placement and generic circumstance of *Stranvaesia*, we obtained a comprehensive taxon sampling of *Stranvaesia* and its allies in the framework of Maleae. All six individuals representing three species recognized in the redefined *Stranvaesia* (Liu et al., 2019) were sampled in this study; they were *S. bodinieri*, *S. nussia*, and *S. oblanceolata*. Because of the ambiguous phylogenetic relationship between *Stranvaesia* and *Photinia*, we also sampled 19 individuals in *Photinia*, representing 16 species out of 20 species currently recognized (Yu and Ku, 1974; Lu and Spongberg, 2003). Additionally, 41 species representing 22 genera have been selected as the outgroup, including 21 genera of Maleae and one genus of Gillenieae (*Gillenia stipulata* (Muhl. ex Willd.) Nutt.). A total of 66 individuals were sampled in our study; 23 of them were sequenced for this study, 21 were from our previous study (Liu et al., 2022), and 22 were downloaded from NCBI. The raw data of these newly sequenced samples were uploaded to Sequence Read Archive (BioProject PRJNA859408). The corresponding accession numbers and voucher information are provided in Table S1

Total genomic DNAs were extracted from silica-gel dried leaves or herbarium specimens using a modified CTAB (mCTAB) method (Li et al., 2013) in the lab of the Institute of Botany, Chinese Academy of Science (IBCAS) in China. The libraries were prepared in the lab of Novogene, Beijing, China using NEBNext^®^ UltraTM II DNA Library Prep Kit, and then paired-end reads of 2 × 150 bp were generated on the NovoSeq 6000 Sequencing System (Novogene, Beijing) with the sequencing depth up to 15.2×.

### 2.2. Single-copy nuclear marker development

SCN marker development followed the pipeline presented in Liu et al. (2021). Briefly, the coding regions of three genomes retrieved from GenBank (*Malus domestica* (Suckow) Borkh., accession number: GCF_002114115.1; *Prunus persica* (L.) Batsch, accession number: GCA_000346465.2; and *Pyrus ussuriensis* Maxim. × *Pyrus communis* L., accession number: GCA_008932095.1) were inputted into MarkerMiner v. 1.0 (Chamala et al., 2015) to identify the putative single-copy genes. The resulting genes were then filtered by successively BLASTing (Altschul et al., 1990, 1997; Camacho et al., 2009) against ten genomes of the family, viz., *Cydonia oblonga* Mill. (accession number: GCA_015708375.1), *Dryas drummondii* Richardson ex Hook. (accession number: GCA_003254865.1), *Fragaria vesca* L. (accession number: GCF_000184155.1), *Geum urbanum* L. (accession number: GCA_900236755.1), *Gillenia trifoliata* (L.) Moench (accession number: GCA_018257905.1), *Malus domestica* (accession number: GCF_002114115.1), *Prunus persica* (accession number: GCF_000346465.2), *Purshia tridentata* (Pursh) DC. (accession number: GCA_003254885.1), *Pyrus ussuriensis*×*P. communis* (accession number: GCA_008932095.1), and *Rosa chinensis* Jacq. (accession number: GCF_002994745.2) using Geneious Prime (Kearse et al., 2012), with the parameters settings in the Megablast program (Morgulis et al., 2008) as a maximum of 60 hits, a maximum E-value of 1 × 10^-10^, a linear gap cost, a word size of 28, and scores of 1 for match and −2 for mismatch in alignments. We first excluded genes with mean coverage > 1.1 for alignments, which indicated potential paralogy and/or the presence of highly repeated elements in the sequences. The remaining alignments were further visually examined to exclude those genes receiving multiple hits with long overlapping but different sequences during BLASTing. It should be noted that the alignments with mean coverage between 1.0 and 1.1 were typically caused by tiny pieces of flanking intron sequences in the alignments. These fragments were considered SCN genes here. The resulting SCN gene reference is available from the Dryad Digital Repository: https://doi.org/10.5061/dryad.hx3ffbghm.

### 2.3. Reads processing and assembly

We processed the raw reads by trimming low-quality bases and sequence adapters using Trimmomatic v. 0.39 (Bolger et al., 2014), with the average quality per base of a four-base sliding window below 15, and reads less than 36 bases removed. Adapters from the deep genome skimming (DGS) and genome resequencing (WGS) data were removed with TruSeq3-PE.fa as input adapter sequences, and those in the transcriptome (RNA-Seq) data were removed with NexteraPE-PE.fa (available from https://github.com/usadellab/Trimmomatic/tree/main/adapters). The trimmed data were then checked with FastQC v. 0.11.9 (Andrews, 2018) to ensure that all adapters were removed and qualified for downstream analysis. The sequencing depth averaged 17.2×, assuming an estimated genome size of around 750 Mb based on *Malus domestica* genome (Table S1).

Plastome assembly followed a two-step strategy proposed by Liu et al. (2019), integrating NOVOPlasty 3.6 (Dierckxsens et al., 2016) and a successive method (Zhang et al., 2015). The latter method combined mapping-based and de novo assembly, and can handle any amount of data to obtain high-quality plastomes. We generated 66 complete chloroplast genomes, 23 of which were newly assembled for this study, and the remaining 43 were downloaded from NCBI. The circularized plastomes were annotated using Geneious Prime (Kearse et al., 2012) with *Photinia prunifolia* (Hook. & Arn.) Lindl. (GenBank accession number MK920279) downloaded from NCBI as the reference for *Photinia* and *Stranvaesia nussia* (GenBank accession number MK920284) for *Stranvaesia*. We then manually checked each coding gene’s start and stop codon in all chloroplast genomes and removed incorrect annotations by translating the sequences into proteins. The final assembled chloroplast genome was converted into the format required by GenBank using GB2sequin (Lehwark and Greiner, 2019), and then submitted to NCBI with the corresponding accession number listed in Table S1.

We utilized the ‘hybpiper assemble’ command of HybPiper v. 2.0.1 (Johnson et al., 2016) to assemble the single-copy nuclear locus of each sample with the parameter “--cov_cutoff 5” based on the SCN gene reference mentioned above. We then summarized and visualized the recovery efficiency using the ‘hybpiper stats’ and ‘hybpiper recovery_heatmap’ commands. Because paralogous genes may impact phylogenetic inference, especially in groups with prevalent reticulate evolution, we performed a paralogous genes search using the post-processing command ‘hybpiper paralog_retriever’ in HybPiper v. 2.0.1 (Johnson et al., 2016) and used the genes without paralog warnings in the subsequent phylogenetic analyses. Due to differences in sequence recovery efficiency among samples because of uneven sequencing coverage in this study, we followed Liu et al. (2022)’s pipeline to further process the assembled SCN genes to remove outlier loci and short sequences, and to account for missing data. Briefly, each SCN gene was aligned by MAFFT v. 7.480 (Nakamura et al., 2018) and clipped by trimAL v. 1.2 (Capella-Gutiérrez et al., 2009) to remove aligned columns with gaps in more than 20% of the sequences and retain sequences with average similarity more than 99.9%. The resulting SCN genes were then concatenated using AMAS v. 1.0 (Borowiec, 2016), and the concatenated genes were then used as input to run Spruceup (Borowiec, 2019) for removing outlier sequences. We also used AMAS v. 1.0 (Borowiec, 2016) to split the processed alignment back into single SCN gene alignments and retrimmed these alignments using trimAL v. 1.2 (Capella-Gutiérrez et al., 2009) with the same parameters as above. Given the potentially limited informativeness in short sequences, we keep the aligned sequences with more than 250 bp length for downstream analysis using a python script (exclude_short_sequences.py, Liu et al., 2022). To remove possible erroneous sequences in the alignments, we used TreeShrink v. 1.3.9 (Mai and Mirarab, 2018) to detect and remove outlier tips with abnormally long branches in each SCN gene tree. The following phylogenetic analysis is based on these shrunk trees and sequences.

### 2.4. Phylogenetic analyses

We inferred the phylogenetic relationships of *Stranvaesia* in the context of Maleae using two sets of data, i.e., nuclear SCN genes and plastid coding sequences (CDSs). In this study, both concatenated and coalescent-based methods were carried out on each data type. All 78 plastid CDSs were extracted from 66 plastomes using Geneious Prime (Kearse et al., 2012). They were aligned by MAFFT v. 7.475 (Nakamura et al., 2018) independently with the “--auto” option and then concatenated by AMAS v. 1.0 (Borowiec, 2016). The best-fit partitioning schemes and/or nucleotide substitution models for downstream analysis were searched using PartitionFinder2 (Stamatakis, 2006; Lanfear et al., 2016), with parameters set to linked branch lengths, Corrected Akaike Information Criterion (AICc) and greedy (Lanfear et al., 2012) algorithm. We first estimated a maximum likelihood (ML) tree with IQ-TREE2 v. 2.1.3 (Minh et al., 2020) with 1000 SH-aLRT and the ultrafast bootstrap replicates, as well as collapsing near zero branches option using the best partitioning schemes and nucleotide substitution models inferred above. An alternative ML tree was inferred using RAxML 8.2.12 (Stamatakis, 2014) with the GTRGAMMA model for each partition and clade support assessed with 200 rapid bootstrap (BS) replicates. Considering possible conflict in evolutionary history among plastid genes, we estimated a coalescent-based species tree based on the 78 plastid CDSs. We inferred individual ML gene tree using RAxML with a GTRGAMMA model, and 200 BS replicates to assess clade support. After collapsing branches with support below 10 using *phyx* (Brown et al., 2017), all 78 gene trees were then used to estimate a species tree using ASTRAL-III (Zhang et al., 2018) with local posterior probabilities (LPP; Sayyari and Mirarab, 2016) to assess clade support. These three trees are available from the Dryad Digital Repository: https://doi.org/10.5061/dryad.hx3ffbghm.

For the nuclear phylogenetic inference, we concatenated all the previously shrunk SCNs sequences with AMAS v. 1.0 (Borowiec, 2016), then used PartitionFinder2 (Stamatakis, 2006; Lanfear et al., 2016) to estimate the best-fit partitioning schemes and/or nucleotide substitution models under the corrected Akaike information criterion (AICc) and linked branch lengths, as well as with the rcluster algorithm (Lanfear et al., 2014). As described above in the plastid phylogenetic analysis, ML trees were estimated by IQ-TREE2 v. 2.1.3 (Minh et al., 2020) with 1000 SH-aLRT using UFBoot2 and collapsing near zero branches option and RAxML 8.2.12 (Stamatakis, 2014) with GTRGAMMA model for each partition and clade support assessed with 200 rapid bootstrap (BS) replicates. Individual ML gene tree were inferred using RAxML with a GTRGAMMA model, and 200 BS replicates to assess clade support. To decrease the systematic error from low-supported clades, we used *phyx* (Brown et al., 2017) to collapse the branches of these shrunk SCN gene trees with support below 10. The processed trees were then used to estimate a coalescent-based species tree with ASTRAL-III (Zhang et al., 2018) using local posterior probabilities (LPP) to assess clade support. These three trees are available from the Dryad Digital Repository: https://doi.org/10.5061/dryad.hx3ffbghm.

### 2.5. Detecting and visualizing nuclear gene tree discordance

Due to the gene tree discordance detected in the inferred phylogeny, we used *phyparts* (Smith et al., 2015) and Quartet Sampling (QS: Pease et al., 2018) to evaluate gene tree conflict in the nuclear datasets. By comparing the gene trees against the ML tree inferred from RAxML, *phyparts* can count the number of concordant and conflicting gene trees at each node of the RAxML tree and yield an Internode Certainty (ICA) score reflecting the degree of conflict. This analysis treated the branch/nodes bootstrap support (BS) of the gene tree inferred from RAxML lower than 50% as uninformative, ignoring such genes. The result of *phyparts* was visualized as pie charts with *phypartspiecharts.py* (by Matt Johnson, available from https://github.com/mossmatters/MJPythonNotebooks/blob/master/phypartspiecharts.py). Based on repeated subsampling of quartets from the input tree and alignment to generate counts of the three possible topologies (and uninformative replicates) and calculate the confidence, consistency, and informativeness of each internal branch, QS analysis can better address phylogenetic discordance with comprehensive and specific information on branch support. Therefore, QS analysis was used to evaluate gene tree conflict with paraments set to 100 replicates and the log-likelihood threshold of 2; this can gather more information on these strongly discordant nodes shown in *phyparts*. The QS result was visualized with plot_QC_ggtree.R (by Shuiyin Liu, available from https://github.com/ShuiyinLIU/QS_visualization).

### 2.6. Network analyses and allopolyploidy analyses

Due to frequent hybridization events and rapid radiation that characterize the Maleae, we used the phylogenomic network method SNaQ, which is implemented in PhyloNetworks (Solís-Lemus et al., 2017), to identify reticulation events, which may explain discordance between the nuclear tree and plastid tree within *Stranvaesia*. SNaQ identifies possible hybridization events and calculates the inheritance probabilities γ and 1-γ representing the transmission of genetic material from two parental lineages to the hybrid. The optimal number of hybridization events is determined based on a pseudolikelihood method. Because of decreasing computational tractability as more species are included, we sampled 12 individuals for use in the SNaQ analysis, consisting of four *Stranvaesia* samples and eight representatives from closely related lineages in Maleae. This sampling strategy covered all potential maternal and paternal lineages of *Stranvaesia*. The quartet concordance factors (CFs) represent the proportion of genes supporting each possible relationship with a given quartet (Larget et al., 2010), and we summarized the CFs based on all 611 nuclear SCN gene trees estimated from RAxML 8.2.12 (Stamatakis, 2014). The species tree estimated with ASTRAL-III (Zhang et al., 2018) was used to generate an optimal starting SNaQ network with no hybridization edges (*h_max_* = 0). The best network (*h_max_* = 0) and the CFs were used to estimate the next optimal network (*h_max_* = 1), and the resulting network was used to estimate the next optimal network (*h_max_* = 2), and so on. We constructed six optimal networks using *h_max_* values ranging from 0 to 5 with 50 independent runs for each *h_max_*. The pseudo-deviance score estimated from each run’s branch lengths and inheritance probabilities can be used to select the optimal phylogenetic network, with lower pseudo-deviance scores indicating a better fit (Solís-Lemus et al., 2017). We plotted *h_max_* (0 to 5) against the log-likelihood score (i.e., network score) of the optimal network for each *h_max_* value to assess the optimal number of hybridization edges. Plotting the decrease in the pseudo-deviance score and observing a leveling-off in the rate of change was used to determine the optimal *h_max_* value.

GRAMPA utilizes the least common ancestor (LCA) mapping algorithm and the representation of multi-labeled trees (MUL-trees) to identify the most probable clade with a polyploid origin. In the absence of polyploidy, a singly-labeled tree would be more parsimonious than any MUL-tree. If a MUL-tree is most parsimonious, the parental lineage(s) involved in a genome doubling event can be inferred from the most parsimonious MUL-tree. Due to frequently reported polyploidy events in some genera in Maleae (Robertson et al., 1991; Liu et al., 2022), we ran the GRAMPA (Thomas et al., 2017) analysis to identify possible allopolyploid and/or autopolyploid scenarios involved in the origin of the *Stranvaesia* clade. We tested if the *Stranvaesia* clade was a result of allopolyploidization (“-h1” inputs), with the remaining nodes investigated as potential secondary parental branches (“-h2” inputs).

### 2.7. Dating analysis and ancestral area reconstruction

We estimated the divergence age and the ancestral areas of *Stranvaesia* in the framework of Maleae using the nuclear SCN datasets. Divergence times were estimated using treePL (Smith and O’Meara, 2012), which uses a penalized likelihood algorithm. The treePL program is suitable for dealing with large datasets with hundreds of taxa by combining two rounds of gradient-based optimization and a partial simulated annealing procedure to achieve speed optimization as well as avoid issues with local optima. We used the best ML tree inferred from RAxML based on the 426 nuclear SCN gene matrix as the phylogenetic backbone for the dating analysis.

A data-driven cross-validation analysis was carried out in treePL (Smith and O’Meara, 2012) to acquire the optimal value for smoothing parameter λ, which determines the appropriate level of rate heterogeneity. We tested 19 smoothing values in multiples of 10 from 1 × 10^−12^ to 1 × 10^6^ and used 1 × 10^−10^ as the best smoothing values for the following dating analysis. Fossils of *Amelanchier peritula* and *A. scudderi* have been discovered in the Florissant Formation, Colorado, USA, and they were dated to Chadronian in Late Eocene (37.2-33.9 million years ago [Ma]). We thus set the stem *Amelanchier* Medik. with a minimum age of 33.9 Ma and a maximum age of 37.2 Ma. The leaf fossil of *Vauquelinia comptoniifolia* from Green River Formation, Colorado, USA has been dated to Eocene (MacGinitie, 1969). We constrained this leaf fossil to 40.4 (the minimum age) and 46.2 Ma (the maximum age). Additionally, a leaf fossil of *Malus* or *Pyrus* from the Republic site, Washington was used to constrain the divergence between *Malus* and *Pyrus* at 46 (the maximum age)-44 (the minimum age) Ma (MacGinitie, 1969). The minimum and maximum ages of the stem of *Gillenia* Moench were constrained to 53.3 and 58.7 Ma, respectively (Zhang et al., 2017). The nuclear phylogeny inferred from RAxML 8.2.12 (Stamatakis, 2014) was applied to construct all dated bootstrap time trees in treePL (Smith and O’Meara, 2012). The resulting dated bootstrap time trees were then used to generate maximum credibility trees in TreeAnnotator v1.10 implemented in BEAST2 (Bouckaert et al., 2014), and the dated best time tree with confidence age intervals was visualized in FigTree v1.4.4.

Biogeographic analyses were conducted using the SCN data as input for BioGeoBEARS v. 1.1.1 (Matzke, 2018) implemented in RASP v. 4.2 (Yu et al., 2015). We delimit five geographical area units according to the distribution of Maleae: (A), East Asia; (B), Europe; (C), Central Asia; (D), North America; (E), South America. The dated best time tree summarized above by TreeAnnotator was used as input to score each taxon to these areas, and the maximum number of areas per node was set to five. We chose the model with the highest AICc_wt value as the best model.

## 3. Results

### 3.1. Single-copy nuclear genes assembly

We filtered a set of 801 SCN genes from thirteen genomes for this phylogenomic study on *Stranvaesia* and its close relatives. The number of genes recovered for each sample varied from 529 (66.0%) to 801 (100%) (Table S2 and Fig. S1). The paralogous genes search using the post-processing command ‘hybpiper paralog_retriever’ in HybPiper v. 2.0.1 (Johnson et al., 2016), identified 367 genes with paralog warnings, which were removed from the analysis, leaving 434 genes for further processing. The number of genes after removing outlier loci, short sequences, and missing data ranged from 232 (53.5%) to 417 (96.1%) (Table S2). Due to the low sequencing coverage and poor SCN genes recovery efficiency of two samples (*Photinia lasiogyna* (Franch.) C.K.Schneid. 3 & 4, Table S1 & S2), they were excluded in our following nuclear phylogenomic analyses.

### 3.2. Plastid phylogenetic relationship and conflict analyses

The aligned plastid supermatrix generated from 78 concatenated CDS sequences of 66 plastomes comprised 68,343 characters, and this data matrix can be accessed from the Dryad Digital Repository: https://doi.org/10.5061/dryad.hx3ffbghm. The phylogenetic trees based on concatenated- and coalescent-based methods yielded almost the same topology (Figs. S2, S3, S4). Therefore, we use the RAxML tree as the plastid phylogeny in subsequent analyses (Figs. 1 & S2). Based on the chloroplast data, *Stranvaesia* was strongly supported to be monophyletic and was sister to a large clade containing *Photinia*, *Sorbus* L., *Cotoneaster* Medik., *Eriobotrya* Lindl., *Rhaphiolepis* Lindl., *Aria* (Pers.) Host, and *Pyrus* L. Within *Stranvaesia*, four individuals of *Photinia lasiogyna* were moderately supported as sister to a clade including two samples of *Stranvaesia bodinieri*; however, this combined clade was sister to a clade containing samples of *Stranvaesia nussia* and *Stranvaesia oblanceolate* with strong support (Figs. 1, S2, S3, S4). Conflict analysis using *phyparts* showed that most gene trees are uninformative with respect to relationships within *Stranvaesia*, and the remaining gene trees were mostly consistent with the topology of the RAxML tree (Fig. S5). The nearly wholly grey pies resulting from limited informative sites demonstrated the limited utility of plastid coding genes to illustrate the degree of discordance in a shallow phylogeny. In contrast, the QS conflict analysis showed strong support (1/-/1; these values quantified the relative support among the three possible resolutions of four taxa and represented the extent of conflict in the node) for almost every node within *Stranvaesia*, implying that no significant topological conflicts existed in these gene trees (Fig. S6).

### 3.3. Nuclear phylogenetic relationship and conflict analysis

The matrix with 426 concatenated and cleaned SCN genes comprised 651,207 bp in aligned length and is available from the Dryad Digital Repository: https://doi.org/10.5061/dryad.hx3ffbghm. Our phylogenetic inference resulted in three nuclear phylogenetic trees based on both concatenated- and coalescent-based methods, and these three trees have nearly similar topologies (Figs. S7, S8, S9). Therefore, we use the RAxML tree as the nuclear topology in the following analyses (Figs. 1 & S7). Our results showed that *Stranvaesia* was recovered as monophyletic in all three nuclear trees, and then sister to a clade containing *Photinia* and *Cotoneaster*. Furthermore, two strongly supported clades within *Stranvaesia* were recognized in the nuclear tree, i.e., the *Weniomeles* clade and the *Stranvaesia* clade (Fig. 1). However, there were some conflicts within *Stranvaesia* between the concatenated-based tree and the species tree. In the ML trees inferred from RAxML (Fig. S7) or IQ-TREE2 (Fig. S8), two samples of *Photinia lasiogyna* were sister to a clade consisting of two individuals of *Stranvaesia oblanceolata* and *Stranvaesia nussia* with moderate support, and then together sister to a clade including samples of *Stranvaesia oblanceolata*. The combined *Stranvaesia* clade was sister to a lineage including two samples of *Stranvaesia bodinieri*. In the species tree, a clade including two samples of *Stranvaesia bodinieri* was sister to the remaining clades. Then, one individual of *Stranvaesia nussia* and a clade consisting of two samples of *Photinia lasiogyna* were successively sister to the clade containing three samples of *Stranvaesia oblanceolata*. The conflict analyses using *phyparts* (Fig. S10) and Quartet Sampling (Fig. S11) were performed to explore the potential gene tree conflicts. The result from *phyparts* showed that most nuclear gene trees supported the sister relationship between the *Weniomeles* clade and the *Stranvaesia* clade, but there were strong gene tree conflicts within the *Stranvaesia* clade (Figs. 1 & S10). Furthermore, the sister relationship between the *Weniomeles* clade and the *Stranvaesia* clade (Figs. 1 & S11) was also confirmed with full support (1/-/1) by the QS analysis.

### 3.4. Network analyses

Considering the possible hybrid origin of *Stranvaesia*, we conducted a SNaQ network analysis. Due to the limitation of computing, only 12 individuals closely related to *Stranvaesia* were included in our study. The optimal number of hybridization events inferred by the SNaQ network analysis was two, as the pseudo-deviance score gradually stabilized when the *h_max_* value exceeded 2 in our pseudo-loglikelihood score plot (Fig. 2b). The optimal network with two hybridization events (Fig. 2a) indicated that *Stranvaesia* resulted from a cross between the lineage of *Stranvaesia bodinieri* (γ = 0.763) and one lineage of a large clade (γ = 0.237), explaining the observed cytonuclear discordance relating to the placement of *Stranvaesia bodinieri* between the plastid tree and the nuclear phylogeny (Fig. 1). The SNaQ network with *h_max_* = 2 showed that the second possible hybrid was *Stranvaesia nussia*, which may have resulted from hybridization between *Stranvaesia lasiogyna* and *Stranvaesia oblanceolate* (Figs. S10 & S11). The other networks with *h_max_* = 3-5 all included hybridization edges similar to the *h_max_* = 2 network (Fig. S12).

**Fig. 2.**
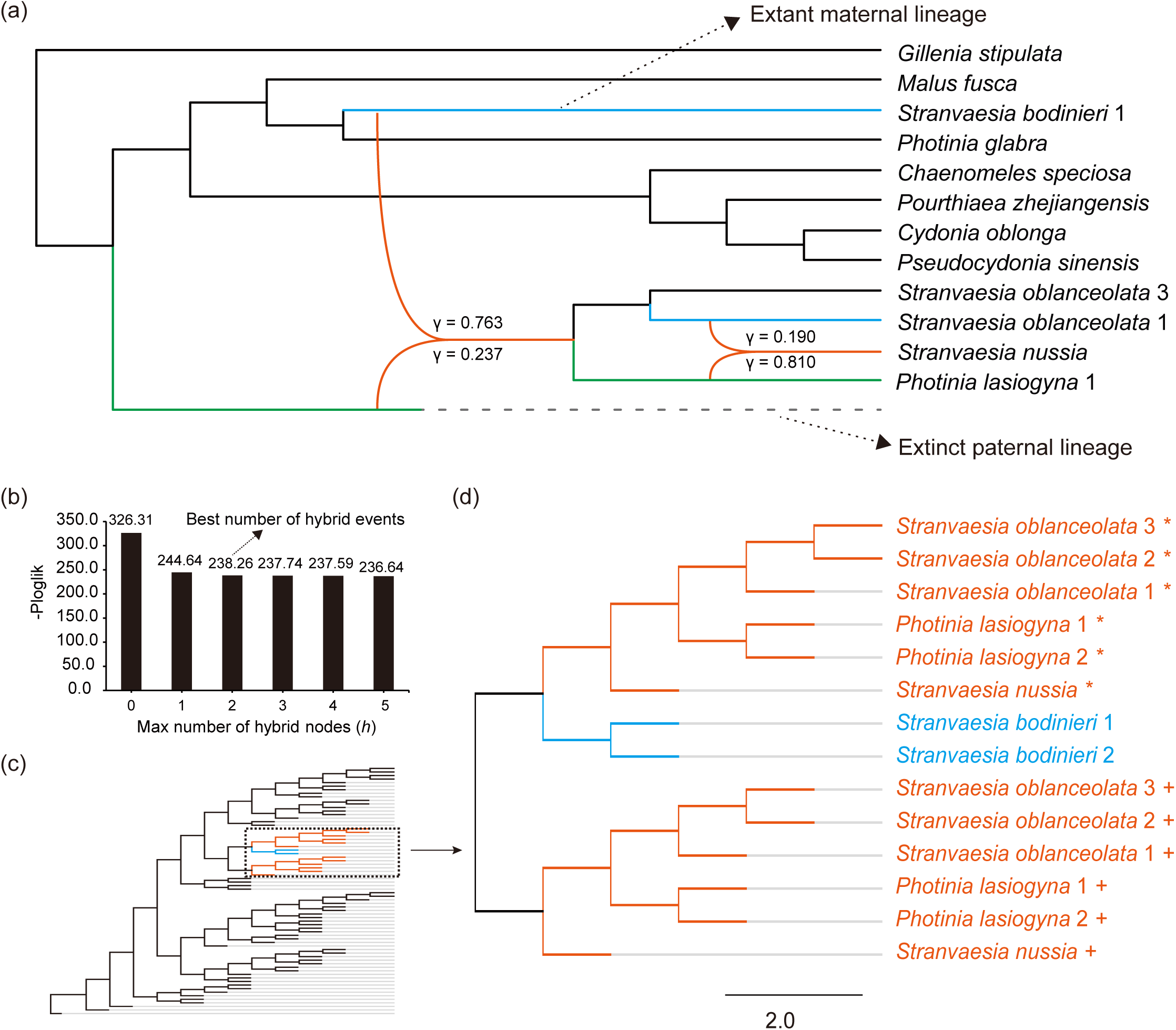
Phylogenetic network analysis from the 12-taxa sampling of *Stranvaesia* and its close allies and GRAMPA allopolyploidy analysis. (a), Species network inferred from SNaQ with a maximum of two reticulations. Orange curved branches indicate the two possible hybridization events with the corresponding inheritance probabilities. Solid lines with different colors denote the extant parental lineages involving hybridization, while dotted lines indicate extinct paternal lineages. (b), The statistics of pseudo-loglikelihood scores (-ploglik) suggest that the optimal network is *h_max_* = 2. (c), Thumbnail of the most parsimonious MUL-trees inferred from GRAMPA analyses based on the nuclear phylogeny with the *Stranvaesia* clade set as a result of allopolyploidization. The figure in detail refers to Fig. S13. (d), The enlarged figure of the part enclosed by the dotted line in c. The clade with multiple labels denotes the polyploid origin, the first tip is indicated by a plus sign and the second tip is shown using an asterisk. One of the *Stranvaesia* clades is sister to the *Weniomeles* clade, and the other *Stranvaesia* clade is sister to the ancestor of *Stranvaesia*, suggesting the allopolyploid origin of the *Stranvaesia* clade.

### 3.5. Multi-labeled trees analysis

Given the potential allopolyploidy events associated with the origin of the *Stranvaesia* clade, we tested an allopolyploidy hypothesis using GRAMPA, with the *Stranvaesia* clade as the possible hybrid and the other taxa in the Maleae as potential parental lineages. The most parsimonious MUL-tree had a lower reconciliation score (score = 62,319) than the estimated singly labeled tree (score = 62,539), implying evidence of polyploidy. The inferred MUL-tree supported that the *Stranvaesia* clade has been an allopolyploid origin between a lineage sister to *Stranvaesia bodinieri* and another lineage sister to the ancestor of *Stranvaesia* (Figs. 2 & S13), which often is considered as another parental lineage that is extinct or not sampled. Despite the discrepancy between the results of the SNaQ network analysis and the GRAMPA analysis, they both recognized *Stranvaesia bodinieri* as one of the possible parental lineages (Figs. 2, S12, S13). However, GRAMPA did not recover another possible parental lineage involved in allopolyploidization, which may have been extinct or not sampled.

### 3.6. Dating analysis and ancestral area reconstruction

The dating analysis showed that the divergence age of the ancestor of the *Stranvaesia* clade and the *Weniomeles* clade was estimated at 41.6 Ma (95% highest posterior density (HPD): 41.22-42.08 Ma) (Figs. 3 & S14) and the ancestral area reconstruction analysis inferred that *Stranvaesia* originated from East Asia (Fig. 3). The stem *Stranvaesia* was estimated at 21.22 Ma (95% HPD: 20.59-22.05 Ma) (Figs. 3 & S14). The crown *Stranvaesia* were estimated at 18.56 Ma (95% HPD: 18.09-19.20 Ma) (Figs. 3 & S14), and *Stranvaesia nussia* originated from ca. 15.13 Ma (95% HPD: 14.21-16.07 Ma).

**Fig. 3.**
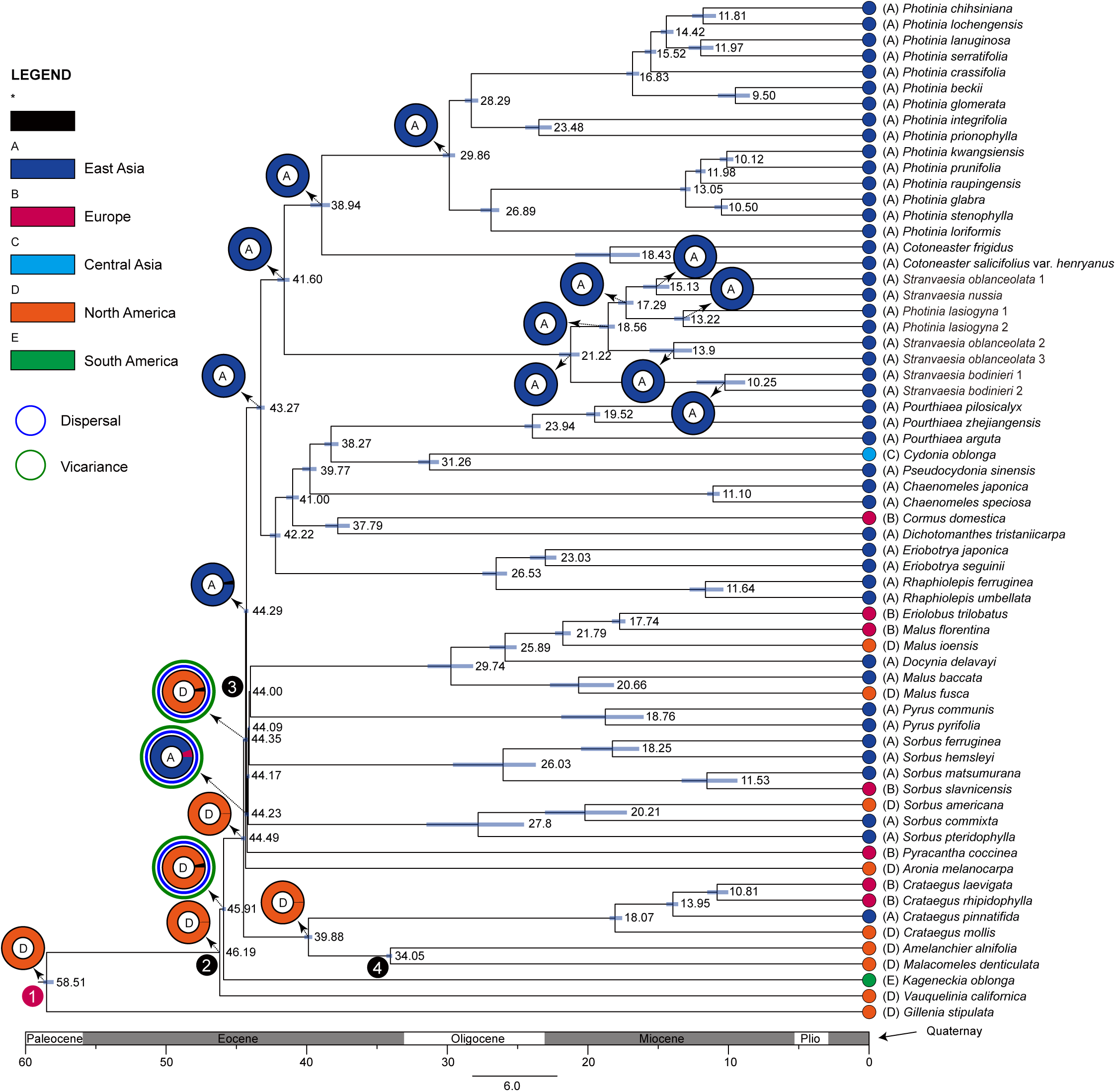
Dated chronogram for the red-fruit genus *Stranvaesia* within Maleae inferred from treePL based on the nuclear data set. Also shown is the ancestral area reconstruction using BioGeoBEARS implemented in RASP, with the colored key identifying extant and possible ancestral ranges. (A), East Asia; (B), Europe; (C), Central Asia; (D), North America; (E), South America. Three fossils, colored black (nodes 2-4), and one divergence time estimate based on previous research, colored purple (node 1), are used as constraints.

## 4. Discussion

The phylogenomic age has made it clear that reticulate evolution, which may originate from either hybridization and/or genome doubling, is a fundamental process in lineage diversification, and accordingly must be considered when modeling the evolution of lineages throughout the Tree of Life. Although analytical approaches have been developed to address hindrances to phylogenetic inference, such as gene tree-species tree incongruence (e.g., ASTRAL, Zhang et al., 2018; *phyparts*, Smith et al., 2015), hybridization (e.g., SNaQ, Solís-Lemus et al., 2017), and genome doubling (e.g., GRAMPA, Thomas et al., 2017), we lack a coherent approach for using phylogenies characterized by reticulate evolution in taxonomic delimitation applications. Using the red-fruit genus *Stranvaesia* as a test case, we demonstrate a workflow for explicitly considering reticulate evolution in the taxon delimitation process. By combining results from cytonuclear conflict, gene tree conflict, network analyses, multiply-labeled trees, morphology, and dating analyses, we identify an entity (*Stranvaesia bodinieri*) that acted as a maternal participant in multiple hybridizations that promoted the early diversification of the genus. Consequently, this entity should be separated from *Stranvaesia* as a new genus. We highlight a broadly-applicable workflow for inferring how analyses of reticulate evolution in phylogenomic data can directly shape taxonomic revisions.

### 4.1. Reticulate evolution challenged the taxonomic circumscription and a practical pipeline for taxonomic treatment

In the past few decades, phylogenomics has revolutionized systematic studies, especially in angiosperm lineages (Leebens-Mack et al., 2019; Li et al., 2019, 2021; Yang et al., 2020; Liu et al., 2021, 2022; Baker et al., 2022). However, phylogenies estimated from hundreds/thousands of nuclear genes and/or plastomes (or mitogenomes in animal studies) often lead to strongly supported phylogenetic inference, although underlying conflict is common despite high support. Hundreds of nuclear genes often result in well-supported bifurcating phylogenetic relationships; however, strongly supported nodes sometimes could not reflect the real evolutionary relationship of taxa, especially in the context of the underlying gene tree conflicts (Smith et al., 2015; Hodel et al., 2021; Liu et al., 2022). Additionally, phylogenetic trees based on different data matrices often result in conflicting relationships (Soltis et al., 1991; Yi et al., 2015; Liu et al., 2017; Liu et al., 2020a). These various topologies may confuse taxonomists and lead to unreasonable taxonomic treatments.

Traditionally, bifurcating phylogenetic trees provided a foundation for taxonomic delimitation, especially when using a criterion of monophyly in the target lineages (Huson and Bryant, 2006; Mindell, 2013). However, mounting evidence shows that reticulate evolutionary histories, such as hybridization, introgression, and polyploidy, promoted the diversity of the angiosperm lineages (Rieseberg and Willis, 2007), and necessitates a new model to reflect complex biological processes better and draw reasonable taxonomic conclusions. Therefore, further analysis will be needed to consider the underlying possible biological processes and evolutionary events. Recently, many programs and approaches have been developed to address reticulate evolutionary histories of taxa by examining underlying processes generating reticulation separately to distinguish the individual effect of separate biological processes (Smith et al., 2015; Solís-Lemus et al., 2017; Thomas et al., 2017; Pease et al., 2018). For example, SNaQ can infer phylogenetic networks with maximum pseudolikelihood under ILS, implementing the statistical inference method (Solís-Lemus and Ané, 2016; Solís-Lemus et al., 2017). HyDe can detect the extent of hybridization using phylogenetic invariants arising under the coalescent model with hybridization (Blischak et al., 2018). Thomas et al. (2017) provided a program GRAMPA, which can distinguish hybridization from polyploid by identifying and placing polyploidy events on a phylogeny.

The genetic age led taxonomists to believe it is not rational to make taxonomic treatments based solely on morphological similarities. Genealogy approaches advocated by Darwinists (Darwinian classification) based on monophyly or common descent have been widely accepted as two additional criteria for biological classification in addition to the overall morphological similarities (referring to the review by Mayr and Bock 2002; but also see Padian 1999). Integrative taxonomy, combining evidence from multiple disciplines, such as morphology, phylogenomics, cytology, and ecology, has been promoted as a standard practice in taxonomic studies (Dayrat, 2005; Schlick-Steiner et al., 2010; Padial et al., 2010). Lineages characterized by reticulate evolutionary events remain a major impediment to taxonomy in our current framework; we cannot make reasonable taxonomic conclusions without carefully explaining the evolutionary processes. In this study, we utilized the red-fruit genus *Stranvaesia* and its allies as a case study to perform a series of phylogenomic analyses, in which we elucidated the important role of *S. bodinieri* in forming *Stranvaesia* and untangled the reticulate evolutionary history of the redefined *Stranvaesia* under significant cytonuclear discord. Therefore, we propose a practical pipeline for making reliable taxonomic treatments when reticulation is suspected. We follow the pipeline for assembling hundreds of SCN genes and plastomes (Liu et al., 2021) and the general procedure for further phylogenetic conflict and network analyses (Liu et al., 2022).

#### Step 1. A comprehensive taxon sampling for the target lineages, including their relatives

Given the potential for reticulation events to be deep or shallow in the evolutionary history of the target group, close or distant relatives of the focal clade may have been involved in its origin. A broad taxon sampling will be helpful for exploring the potential paternal and maternal parents.

#### Step 2. Accurate phylogenetic inference

Two distinctly inherited datasets, hundreds/thousands of nuclear SCN genes and plastid (in plant studies) and/or mitochondrial (in aminal studies) coding genes, are needed to assess phylogenomic discordance and conduct historical biogeographic analyses. Based on the size of data sets and the targeted lineages, paralogous loci could be discarded directly (e.g., Crowl et al., 2019; Bagley et al., 2020) or estimated by the tree-based orthology inference (Yang and Smith, 2014; Morales-Briones et al., 2022). Additionally, given the sample’s uneven and limited sequencing coverage, outlier sites, missing data, and short sequences in the assembled SCN genes need to be trimmed using several programs, such as trimAL, Spruceup, and TreeShrink. Generate nested datasets based on the number of genes and samples and then perform the phylogenetic analysis using concatenated supermatrix (RAxML, IQ-TREE2, and MrBayes) and coalescent-based methods (ASTRAL-III, SVDquartets, MP-EST, Quartet MaxCut, etc.).

#### Step 3. Phylogenomic conflict analyses

Compare the plastid (or mitochondrial, in the case of animals) and nuclear topologies. Detect and visualize gene tree discordance using *phyparts* and/or Quartet Sampling. Assessing cytonuclear discord may suggest hypotheses to test using additional analyses in **Step 4**.

#### Step 4. Phylogenetic network and historical biogeographic analyses

Observed phylogenomic discordance may be due to ILS, allopolyploidy, and/or hybridization. Coalescent simulations could adequately measure the coalescent model’s goodness-of-fit with ILS explaining the gene tree discordance. When ILS cannot explain discordance sufficiently, phylogenetic network analyses are appropriate for testing hybridization, while limiting the number of terminals (not more than 30) for evaluation. If the number of tips exceeds 30, subsampling the species for phylogenetic network analysis is necessary. Subsequent network analyses using different datasets at various taxonomic levels may be adopted as an alternative strategy to circumvent the limitation on the number of species. Multiply-Labeled Tree Analysis (GRAMPA) and/or chromosome data could test the role of allopolyploidization and/or autopolyploidization. Historical biogeographic analyses, including fossil and living species, will accurately provide the divergence time and area in which the potential evolutionary events occurred.

#### Step 5. Taxonomic treatments

Propose appropriate taxonomic treatments integrating evidence from multiple disciplines, such as morphology, phylogenomics, cytology, biogeography, and ecology. Considering the prevalent reticulations in some lineages, we recommend that infrageneric hybrids be treated in the same genus. If the parents are distantly related and do not share many similarities, it would be justified to consider the hybrids to be of the same status as the parents.

### 4.2. Allopolyploidy and introgression promoted the origin and diversification of Stranvaesia

Reticulate evolution has greatly challenged accurate phylogenetic inference and corresponding taxonomic treatments. Efforts to resolve the generic circumscription of *Photinia* and *Stranvaesia* have been made in the past decades (Guo et al., 2011; Lo and Donoghue, 2012; Liu et al., 2019, 2022); however, significant controversies arose due to extensive reticulate evolution, such as frequent hybridization and allopolyploidy events. Elucidating the evolutionary history of *Stranvaesia* and its allies will provide insights into their generic delimitation. We used hundreds of nuclear SCN genes and plastomes in this study to clarify the complex relationship between *Photinia* and *Stranvaesia* with broad sampling. Our phylogenomic analyses inferred from these two datasets provided a strongly supported phylogenetic backbone of *Stranvaesia* in the framework of Maleae. Cytonuclear discordance was detected between the chloroplast and nuclear gene trees; further gene tree discordance analyses (e.g., *phyparts* and QS) revealed that the varied phylogenetic position of *Stranvaesia bodinieri* did not solely result from ILS. Furthermore, based on the results from the phylogenetic network analysis and MUL-trees analysis, *Stranvaesia bodinieri* may have acted as one of the maternal lineages involved in multiple hybridization events, promoting the origin of the *Stranvaesia* clade. Therefore, *Stranvaesia bodinieri* should be separated from *Stranvaesia* as a new genus, *Weniomeles*. We also discuss the taxonomic delimitation challenged by reticulate evolution events and made nomenclatural treatment. Below we contextualize and discuss the details of our results.

Although the monophyly of *Stranvaesia* was recovered in our nuclear and plastid phylogenies with strong support, cytonuclear discordance was detected within this genus. *Stranvaesia bodinieri* was either nested in the *Stranvaesia* clade in plastid topology (Figs. 1, S2, S3, S4) or formed a sister relationship with a lineage containing the remaining *Stranvaesia* species in the nuclear tree (Figs. 1, S7, S8, S9). This conflict may have resulted from several potential processes, such as ILS and gene flow (hybridization, introgression, and allopolyploidy) (Rieseberg and Soltis, 1991). Because discordance between gene trees in each dataset may contribute to cytonuclear discordance, we separately performed conflict analyses on nuclear- and chloroplast-inferred phylogenies. The *phyparts* results from the nuclear phylogeny showed that 271 SCN genes (63.6%) supported the sister relationship between the *Stranvaesia bodinieri* clade and the *Stranvaesia* clade, contrasting with the 37 unsupported SCN genes (8.7%) (Figs. 1 & S10). However, all sampled quartet replicates in the QS analysis supported this node (QC = 1), with all trees informative when likelihood cutoffs are used (QI = 1) (Figs. 1 & S11). Our further phylogenetic network analysis showed that ILS (*h_max_ =* 0) could not fully explain this conflict (Fig. S12). Therefore, the most likely source of the conflict placement of *Stranvaesia bodinieri* is gene flow rather than ILS, especially in the context of the frequent hybridization events of Maleae (Robertson et al., 1991; Lo and Donoghue, 2012; Liu et al., 2019, 2020a, 2020b, 2022).

The inferred optimal network with two possible hybridization events demonstrated that the *Stranvaesia* clade in this study possibly originated from hybridization between the ancestor of *Stranvaesia bodinieri* (γ = 0.763) and the ancestor of a large clade (γ = 0.237), including *Chaenomeles* Lindl., *Cydonia* Mill., *Malus* Mill., *Photinia* Lindl., *Pourthiaea* Decne., *Pseudocydonia* (C.K.Schneid.) C.K.Schneid., and *Stranvaesia* (Fig. 2). This uneven proportion could be interpreted as introgression (Solís-Lemus et al., 2017), i.e., the repeated backcrossing between the hybrid and the ancestor of *Stranvaesia bodinieri*. Introgression between different genera leads to speciation and diversification has also been observed in other taxa, such as Fagaceae (Zhou et al., 2022). Polyploidy events are prevalent in plants, and at least 15% of speciation events in angiosperms are estimated to be driven by polyploidization (Wood et al., 2009; Mayrose et al., 2011). Given the potential allopolyploidy events between these two different genera, we tested the possible origin of allopolyploidization for the newly defined *Stranvaesia* using GRAMPA (Thomas et al., 2017). The most parsimonious MUL-tree (Figs. 2 & S13), which showed a lower reconciliation score than a singly-labeled tree, supported the allopolyploidy origin of the newly defined *Stranvaesia*. The ancestor of *Stranvaesia bodinieri* may have been the maternal parent, and the paternal parent may have been extinct (Fig. 4). This hypothesis is consistent with the SNaQ result (Fig. 2). Recently, *Malus sikkimensis* (Wenz.) Koehne has been hypothesized to have originated via allopolyploidization based on phylogenomic and chromosome evidence (Liu et al., 2022), and the ploidy level varies from diploid to tetraploid (Liang, 1986, 1987, 1997; Liang and Li, 1993; Liang et al., 1996). However, as the only recorded chromosome count in *Stranvaesia*, *Stranvaesia glaucescens* Lindl. (= *Stranvaesia nussia* (D.Don) Decne.) has been registered as diploid with *x* = 17 (Mehra et al., 1973; Singhal et al., 1990). Therefore, we speculated that diploidization may have occurred following the initial allopolyploidization, and the following frequent backcrosses between the hybrid (the ancestor of *Stranvaesia*) and the ancestor of *S. bodinieri* resulted in the uneven inheritance of genetic material. Our dating and ancestral area reconstruction analyses (Figs. 3 & S14) showed that allopolyploidization events may have occurred in the middle Miocene in East Asia, coinciding with the climatic cooling event after the Middle Miocene Climatic Optimum (MMCO, around 15 Mya) (Flower and Kennett, 1994). The paleoclimatic events might have significantly impacted the speciation and diversification of *Stranvaesia* and its allies.

**Fig. 4.**
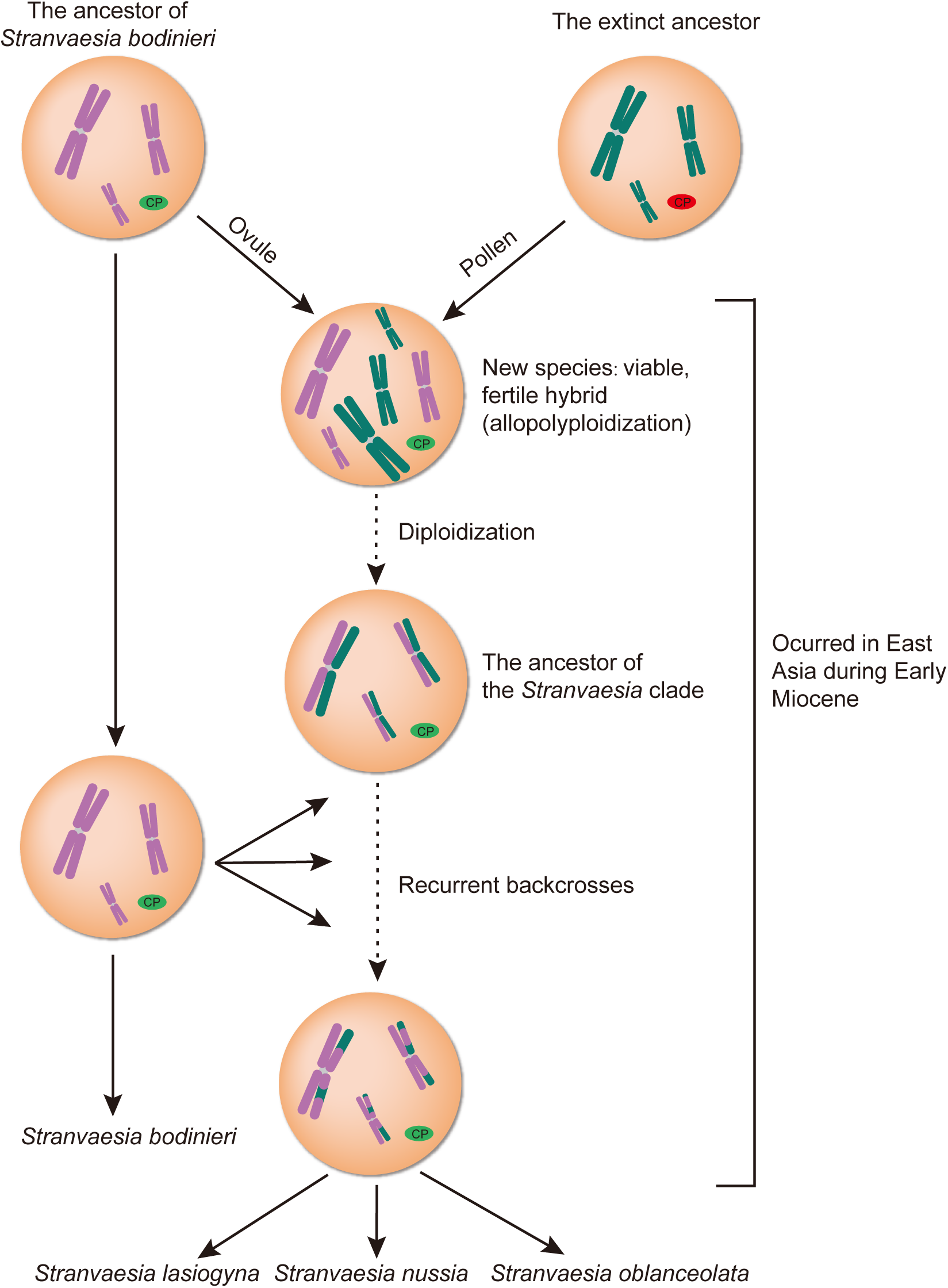
A hypothetical scenario of the origin of *Stranvaesia*. cp, chloroplast genome.

### 4.3. Phylogenetic and taxonomic implications for Photinia and Stranvaesia

The placement of *Stranvaesia* has been a long-standing taxonomic challenge due to its resemblance to its close relatives. Morphological traits used to distinguish *Stranvaesia* from its related genera have been controversial and have been updated over time (Roemer, 1847; Decaisne, 1874; Wenzig, 1883; Focke, 1888; Koehne, 1890, 1891; Rehder, 1940, 1949; Yu, 1974; Lu et al., 1990, 1991; Li, et al. 1992; Zhang, 1992; Lu and Spongberg, 2003; Guo et al., 2011, 2020; Liu et al., 2019). Our phylogenomic analyses, which integrated data from biparentally inherited nuclear genes (426 SCN genes) and maternally inherited plastid coding sequences (78 CDSs), using concatenated or coalescent-based methods, have well resolved the generic circumscription of *Photinia* and provided solid evidence for transferring *Photinia lasiogyna* to *Stranvaesia* (Figs. 1, S2, S3, S4, S7, S8, S9). Due to shared morphological characteristics, including pollen traits of some *Photinia* species (Yu, 1974; Lu and Spongberg, 2003; Pathak et al., 2019), *Photinia lasiogyna*, an endemic species of China, has been traditionally classified in the genus *Photinia* (Yu, 1974; Lu and Spongberg, 2003; Guo, et al. 2020), albeit rarely in *Eriobotrya* (Franchet and Delavay, 1890) and *Pyrus* (Christenhusz et al., 2018). A recent phylogenetic study based on limited nuclear and/or plastid markers also recovered a close relationship between *Photinia lasiogyna* and other *Photinia* species (Guo et al., 2020), in contrast to our phylogenomic result (Figs. 1, S2, S3, S4, S7, S8, S9). However, this topological conflict may have resulted from the limited informative sites of the nuclear (*PepC*) and chloroplast DNA regions (*trnS-trnG*, *psbA-trnH*, and *trnL-trnH*) used in Guo et al. (2020)’s study, given the low resolution of the *Photinia* clade recovered.

Our phylogenetic network analysis also provided insights into the potential hybrid origin of *Stranvaesia nussia*, possibly resulting from a cross between *Photinia lasiogyna* (γ = 0.81) and *S. oblanceolata* (γ = 0.19), with the former acting as the paternal parent and the latter as the maternal parent (Figs. 2 & S12). However, this study did not aim to elucidate the evolutionary history of *Stranvaesia nussia* fully. We hope to further test its hybrid origin with phylogenomic evidence through population-level sampling.

*Stranvaesia* has been redefined based on morphological and phylogenomic evidence, characterized by a cluster of sclereids between locules in the flesh of pomes (Kalkman, 1973; Liu et al., 2019). Upon careful examination of specimens of *Photinia lasiogyna* in the herbarium PE, we found that this species also possesses a cluster of sclereids between locules in the flesh of pomes. We, herein, formally transferred *Photinia lasiogyna* and its variety to *Stranvaesia* as below.

***Stranvaesia lasiogyna*** (Franch.) B.B.Liu, **comb. nov.**

≡ *Eriobotrya lasiogyna* Franch., Pl. Delavay. 225. 1890. ≡ *Photinia lasiogyna* (Franch.) C.K.Schneid., Repert. Spec. Nov. Regni Veg. 3: 153. 1906. ≡ *Pyrus avalon* M.F.Fay & Christenh., Global Fl. 4: 96. 2018. Type: China. Yunnan, in silvis montanis ad fauces San-tchang-kiou supra Hokin, alt. 2300 m., 22 May 1884, *J.M. Delavay 732* (lectotype, designated by Idrees et al. (2022: 31): P [barcode P02143141]!; isolectotypes: P [barcode P02143142]!, US [barcode 00097489]!, image A [barcode 00026747]! with plant material from P02143141).

= *Stranvaesia glaucescens* var. *yunnanensis* Franch., Pl. Delavay. 226. 1890. Type: China. Yunnan, in silvis supra Che-tong, prope Tapin-tze, May 18, 1885, *J.M. Delavay 1992* (lectotype, designated by Idrees et al. (2022: 31): P [barcode P02143161]!; isolectotype: P [barcode P02143140]!).

= *Photinia mairei* H.Lév., Bull. Acad. Int. Géogr. Bot. 17: 28. 1916. Type: China. rochers-brousse des mont a Kiao-me-ti, May 1911-1913, *E.E. Maire s.n.* (holotype: E [barcode E00011316]!; isotype: A [barcode 00038571]!).

Distribution: China (Sichuan and Yunnan).

***Stranvaesia lasiogyna*** var. ***glabrescens*** (L.T.Lu & C.L.Li) B.B.Liu, **comb. nov.**

≡ *Photinia lasiogyna* var. *glabrescens* L.T.Lu & C.L.Li, Acta Phytotax. Sin. 38(3): 278. 2000. Type: China. Jiangxi, Shangrao, 4 May 1972, *Jiangxi Exped. 1071* (holotype: PE [barcode 00336583]!; isotype: PE [barcode 00336582]!).

Distribution: China (Fujian, Guangdong, Guangxi, Hunan, Jiangxi, Sichuan, Yunnan, and Zhejiang).

### 4.4. A new genus, Weniomeles: evidence from morphological and phylogenomic data

The phylogenetic trees inferred from plastomes and multiple nuclear loci datasets supported the monophyly of the *Stranvaesia* sensu Liu et al. (2019), including *S. bodinieri*. However, the phylogenetic placement of *S. bodinieri* varied greatly, either embedded in (plastid tree, Figs. 1, S2, S3, S4) or sister to (nuclear tree, Figs. 1, S7, S8, S9) the *Stranvaesia* clade. The nrDNA results presented an alternative topology with nesting in the redefined *Stranvaesia*, which was very similar to the plastid tree (Liu et al., 2019). This conflict may be explained by the incomplete concerted evolution of nrDNA (Weitemier et al., 2015; Fonseca and Lohmann, 2020) and gene tree discordance between nrDNA and the other nuclear genes. Guo et al. (2020) also indicated the close relationship between *Stranvaesia bodinieri* and the redefined *Stranvaesia* based on nuclear *PepC* and chloroplast data. In this study, phylogenomic discordance and network analyses indicated that *Stranvaesia bodinieri* was involved in the origin of the *Stranvaesia* clade as the maternal parent, followed by recurrent backcrosses (Figs. 1 & 2). Intergeneric hybridization is prevalent in Maleae (Robertson et al., 1991) and some genera such as *Micromeles* Decne., *Phippsiomeles* B.B.Liu & J.Wen, and *Pseudocydonia* have been inferred to have a hybrid origin (Lo and Donoghue, 2012; Liu et al., 2019). Based on the golden criterion (i.e., monophyly) in classification, it will be reasonable to classify *Stranvaesia bodinieri* (the *Weniomeles* clade) and the *Stranvaesia* clade to be one genus or two genera. However, evolutionary (Darwinian) and cladistic (Hennigian) classifications both request the consideration of common descent (Hörandl, 2006). *Stranvaesia bodinieri* and the *Stranvaesia* clade do not have a common ancestor, and the latter originated from allopolyploidization between the former and an extinct clade. We thus propose the *Stranvaesia bodinieri* clade to be a new genus *Weniomeles* rather than a member of *Stranvaesia*.

Morphologically, *Weniomeles bodinieri* is distinct from *Stranvaesia nussia* and *S. oblanceolata* in the number of ovaries, the length of petioles, and the indumentum of rachis, pedicels and hypanthium (Guo et al., 2020). Additionality, this new genus *Weniomeles* has thorns on the stems and/or branches (Fig. 5F, Guo et al., 2011), contrasting to the absence in the *Stranvaesia* clade. Furthermore, the leaf epidermis analysis conducted by Guo et al. (2011) did not detect the same epidermis structure between *Photinia davidsoniae* (= *Stranvaesia bodinieri*) and *Photinia nussia* (= *Stranvaesia nussia*).

**Fig. 5.**
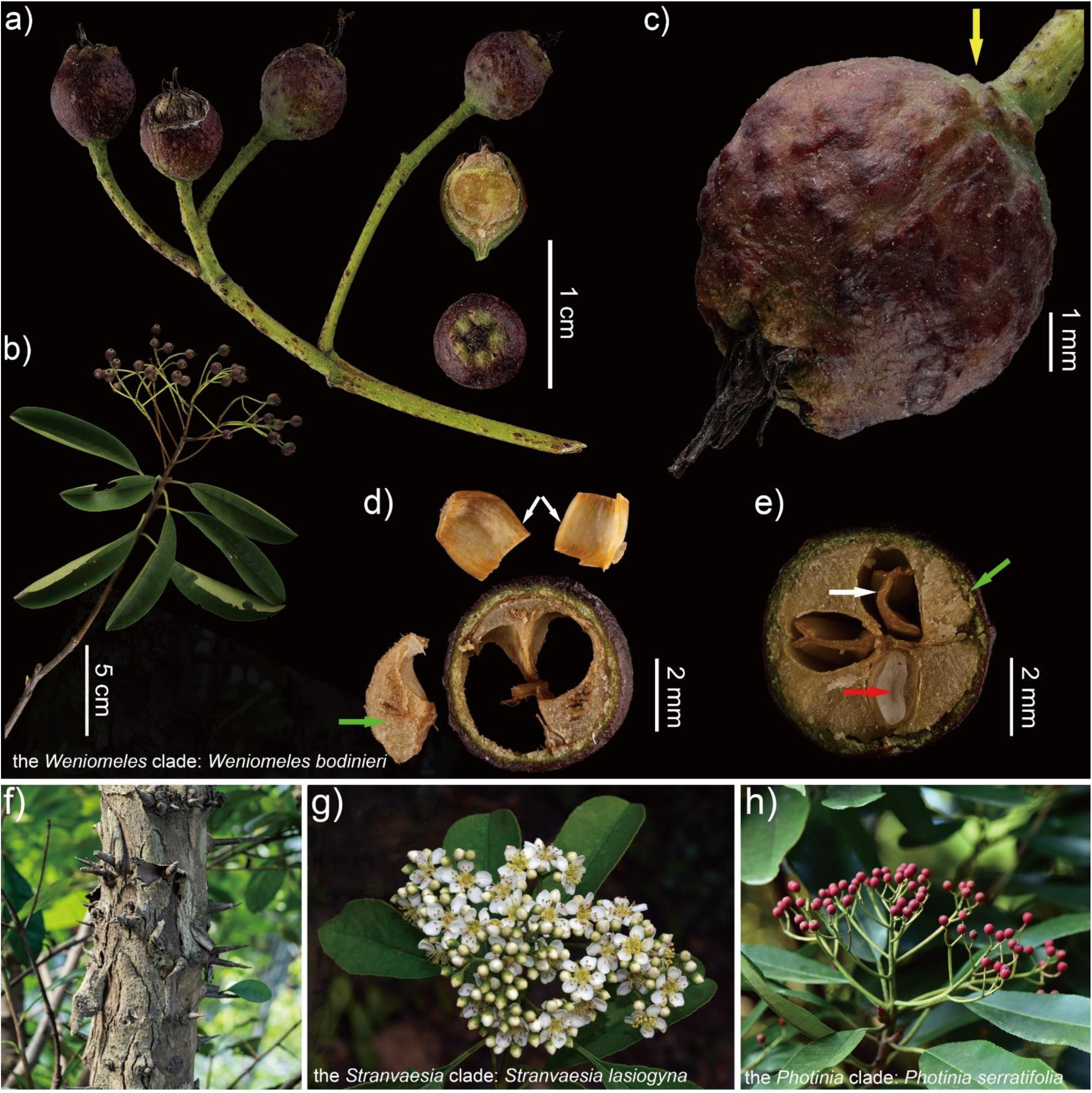
Fine structure and morphological characteristics of the represented clades. a)-f), the *Weniomeles* clade (*Weniomeles bodinieri*): Bin-Jie Ge; a)-b), infructescence; c), fruit (yellow arrowhead); d)-e), cross-section of fruit, showing the fruit core with multilocular separated by a layer of sclereids (red and white arrowhead) and a cluster of sclereids at the top of the locules (green arrowhead); f) thorns on the stems. g), the *Stranvaesia* clade (*Stranvaesia lasiogyna*): Long-Yuan Wang. h), the *Photinia* clade (*Photinia serratifolia*): Xin-Xin Zhu.

Below, we formally describe the new genus *Weniomeles* and make the nomenclatural transfer.

***Weniomeles*** B.B.Liu, **gen. nov.** Type: ***Weniomeles bodinieri*** (H.Lév.) B.B.Liu ≡ *Photinia bodinieri H.Lév*.

*Diagnosis*. *Weniomeles*, characterized by purple-black fruits (Fig. 5A-C), trunk and/or branches with thorns (Fig. 5F), and fruit core with multilocular (Fig. 5E red and white arrowhead) separated by a layer of sclereids and a cluster of sclereids at the top of the locules (Fig. 5D green arrowhead), could be easily distinguished from its close allies, *Stranvaesia* (Fig. 5G).

*Description*. Evergreen trees, usually 6-25 m tall, with a trunk up to 1.4 m in diameter, usually with thorn branches. Unregularly peeling bark, gray-brown when young, brown when old, armed. Petiole (0.8-) 1-1.5 cm, glabrescens; leaf blade oblong, elliptic or obovate to oblanceolate or narrowly lanceolate, 5-10 (−15) × (1.5-) 2-5 cm, veins 10-16 (−20) pairs, both surfaces glabrous or initially slightly pubescent along veins, glabrescent, base cuneate, margin serrate, apex acute to acuminate, obtuse, rarely concave. Compound corymbs terminal, compact, 5-8 × 5-10 cm, many flowered; rachis and pedicels appressed pubescent; bracts caducous, lanceolate or linear, 2-4 mm, pubescent. Pedicel 4-8 mm. Flowers 1-1.5 cm in diameter. Hypanthium cupular, abaxially glabrous to sparsely appressed pubescent. Sepals broadly triangular, 1-2 mm, apex acute or obtuse. Petals white, ovate, ellipsoidal, suborbicular, 5-6 mm long, 3.5-4 wide, glabrous, shortly clawed, apex obtuse or emarginate, base pubescent. Stamens 20, shorter than petals. Styles 2 or 3, connate from base to middle, white villous basally; ovary 2-3-loculed. Fruit purple-black, globose or ovoid, 7-10 mm in diam., glabrous; seeds usually 2, rarely 3, 4, brown, ovoid, 4-5 mm.

*Distribution*. China (Anhui, Fujian, Guangdong, Guangxi, Guizhou, Hubei, Hunan, Jiangsu, Shaanxi, Sichuan, Yunnan, and Zhejiang), Indonesia, and North Vietnam.

*Etymology*. This new genus is named in honor of Prof. Jun Wen (National Museum of Natural History, Smithsonian Institution) for her significant contributions to bridging the Sino-American plant systematic community.

***Weniomeles bodinieri*** (H.Lév.) B.B.Liu, **comb. nov.**

≡ *Photinia bodinieri* H.Lév., Repert. Spec. Nov. Regni Veg. 4: 334. 1907. ≡ *Pyrus eureka* M.F.Fay & Christenh. Global Fl. 4:103. 2018. ≡ *Stranvaesia bodinieri* (H.Lév.) B.B.Liu & J.Wen, J. Syst. Evol. 57(6): 686. 2019. ≡ *Stranvaesia bodinieri* (H.Lév.) Long Y.Wang, W.B.Liao & W.Guo, Phytotaxa 447(2): 110. 2020, isonym. Type: CHINA, Kouy-Tchéou (now Guizhou): environs de Kouy-Yang, mont. du Collège, ca et là autour des villages, 18 May 1898, *E. Bodinier 2256* (lectotype, designated by Liu et al. (2019: 686): P [barcode P02143207]!, isolectotypes: A [barcode 00045584]!, E [barcode E00010998]!, P [barcode P02143208]!, P [barcode P02143209]!).

= *Photinia davidsoniae* Rehder & E.H.Wilson, Pl. Wilson. 1: 185. 1912 ≡ *Pyrus davidsoniae* (Rehder & E.H.Wilson) M.F.Fay & Christenh. Global Fl. 4: 101. 2018. Type: CHINA, Western Hupeh (Hubei): near Ichang (Yichang), alt. 300‒600 m., April 1907, *E.H. Wilson 685* (lectotype, selected by Vidal (1968), first step “type”; second step, designated by Liu et al. (2019: 687): A [barcode 00038567]! excl. the fruits and seeds in the packet, isolectotypes: BM [barcode BM000602130]!, E [barcode E00011306]! excl. the fruiting branch, GH [barcode 00045598]! excl. the fruiting branch, HBG [barcode HBG511078]! excl. the fruiting branch, US [barcode 00097494]! excl. the fruiting branch). (the detailed type information refers to Liu et al. (2019)).

= *Hiptage esquirolii* H.Lév. Repert. Spec. Nov. Regni Veg. 10:372. 1912. Type: CHINA, Kouy-Tchéou (now as Guizhou): Choui -Teou, route de Tin-Pan-Lo-Fou, alt. 900 m, 4 May 1900, *J Esquirol 2097* (lectotype, designated by Liu et al. (2019: 687): E [barcode E00011307]!, isolectotypes: A [barcode 00015103]!, A [barcode 00045102]!).

Distribution: China (Anhui, Fujian, Guangdong, Guangxi, Guizhou, Hubei, Hunan, Jiangsu, Shaanxi, Sichuan, Yunnan, and Zhejiang), Indonesia, and Vietnam.

***Weniomeles bodinieri*** (H.Lév.) B.B.Liu var. *longifolia* (Cardot) B.B.Liu, **comb. nov.**

≡ *Photinia bodinieri* H.Lév. var. *longifolia* Cardot, Notul. Syst. (Paris) 3: 374. 1918. ≡ *Stranvaesia bodinieri* var. *longifolia* (Cardot) B.B.Liu & J.Wen, J. Syst. Evol. 57(6): 687. 2019. Type: CHINA, Kouei Tchéou (now as Guizhou Province): grande route Kouei Tchéou au Kuangsi (Guangxi Province), Kout’ong (now as Gudong Xiang, Pingtang County), 22 May 1899, *Beauvais J. 175* (lectotype, designated by Liu et al. (2019: 687): P [barcode P02143211]!, isolectotype: P [barcode P02143210]!).

Distribution: China (Guizhou).

***Weniomeles bodinieri*** (H.Lév.) B.B.Liu var. ***ambigua*** (Cardot) B.B.Liu, **comb. nov.**

≡ *Photinia davidsoniae* Rehder & E.H.Wilson var. *ambigua* Cardot, Notul. Syst. (Paris) 3: 374. 1918. Type: CHINA, Su-Tchuen (Sichuan): Eul Se Yug, vallée du Yalory, alt. 2000 m, 5 May 1911, *Legendre 834* (**lectotype, designated here**: P [barcode P02143164]!; isolectotype: P [barcode P02143165]!).

Distribution: China (Sichuan).

***Weniomeles bodinieri*** (H.Lév.) B.B.Liu var. ***pungens*** (Cardot) B.B.Liu, **comb. nov.**

≡ *Photinia davidsoniae* Rehder & E.H.Wilson var. *pungens* Cardot, Notul. Syst. (Paris) 3: 374. 1918. Type: CHINA, Hubei: Ichang, *A. Henry 7174* (holotype: P [barcode P02143163]!).

Distribution: China (Hubei).

***Weniomeles atropurpurea*** (P.L.Chiu ex Z.H.Chen & X.F.Jin) B.B.Liu, **comb. nov.**

≡ *Photinia atropurpurea* P.L.Chiu ex Z.H.Chen & X.F.Jin, J. Hangzhou Univ., Nat. Sci. Ed. 20(4): 393. 2021. Type: CHINA, Zhejiang: Taishun, Zuoxi, Lishuqiu, alt. 400 m, 3 May 2020, *Z.H. Chen, Z.P. Lei & W.Y. Xie TS20050316* (holotype: ZM; isotype: ZM).

Distribution: China (Zhejiang).

## 5. Conclusions

We developed and demonstrated the utility of a pipeline for explicitly incorporating reticulation into taxonomic treatments, using results quantifying reticulate evolution from phylogenomic datasets. Our results resolved the placement of *Stranvaesia* in the framework of Maleae. All six phylogenetic trees from coalescent- and concatenated-based methods based on nuclear and plastid data support a monophyletic *Stranvaesia*, including *Photinia lasiogyna*. Extensive gene tree conflicts among nuclear gene trees suggest a complex evolutionary history of redefined *Stranvaesia*, in which ILS, hybridization, and allopolyploidy may have been involved in its diversification. The detected cytonuclear discordance of the *Stranvaesia bodinieri* clade can be explained by the allopolyploidization and the subsequent recurrent backcrosses. Allopolyploidy and introgression may have been involved in the origin of the redefined *Stranvaesia*, in which the ancestor of *Stranvaesia bodinieri* may have acted as the maternal parent and an extinct lineage as the paternal parent. We proposed the descendant of the maternal parent (*Stanvaesia bodinieri*) of the redefined *Stranvaesia* as a new genus, *Weniomeles*, characterized by purple-black fruits, trunk and/or branches with thorns, and fruit core with multilocular separated by a layer of sclereids and a cluster of sclereids at the top of the locules. Given the extensive reticulation in *Stranvaesia* and its allies, this lineage represents a good case study to untangle the reticulate evolution scenario using phylogenomic analyses and the following taxonomic circumscription. With an increasing number of reported complex histories of plant lineages based on increased amounts of sequence data and continuously improved analytical approaches, this phylogenomic case study of *Stranvaesia* implies that taxonomic delimitation must consider evolutionary results from many data types.

## CRediT authorship contribution statement

**Ze-Tao Jin**: Methodology, Software, Investigation, Formal analysis, Writing – original draft. **Dai-Kun Ma**: Methodology, Software, Formal analysis, Writing – original draft. **Richard G.J. Hodel**: Methodology, Software, Writing – review & editing. **Hui Wang**: Investigation, Formal analysis. **Guang-Ning Liu**: Writing – review & editing. **Chen Ren**: Software, Writing – review & editing. **Bin-Jie Ge**: Resources, Formal analysis. **Qiang Fan**: Resources. **Shui-Hu Jin**: Writing – review & editing. **Chao Xu**: Writing – review & editing, Investigation. **Jun Wu**: Writing – review & editing. **Bin-Bin Liu**: Conceptualization, Methodology, Writing – review & editing, Resources, Supervision.

## Data availability

The raw sequence data were deposited in the NCBI Sequence Read Archive (SRA) database under the BioProject PRJNA859408. Alignments and gene trees of all datasets used in this study are available from the Dryad Digital Repository: https://doi.org/10.5061/dryad.hx3ffbghm.

## Acknowledgements

The phylogenomic analyses have been run on the Dell T7920 workstation (owned by Bin-Bin Liu, Institute of Botany, Chinese Academy of Sciences), and the Bioinformatics Center of Nanjing Agricultural University provides support for the analysis. This work was supported by National Natural Science Foundation of China (grant number 32000163 to Bin-Bin Liu and 32270216 to Bin-Bin Liu); the Youth Innovation Promotion Association CAS (grant number 2023086 to Bin-Bin Liu); Project of National Plant Specimen Resource Center (NPSRC) (grant number E0117G1001); and National Wild Plant Germplasm Resource Center for Shanghai Chenshan Botanical Garden (grant number ZWGX2102).

## Appendix A. Supplementary data

Supplementary data to this article can be found online.

## References

Altschul, S.F., Gish, W., Miller, W., Myers, E.W., Lipman, D.J., 1990. Basic local alignment search tool. J. Mol. Biol. 215, 403–410. https://doi.org/10.1016/S0022-2836(05)80360-2.

Altschul, S.F., Madden, T.L., Schaffer, A.A., Zhang, J.H., Zhang, Z., Miller, W., Lipman, D.J., 1997. Gapped BLAST and PSI-BLAST: a new generation of protein database search programs. Nucleic Acids Res. 25, 3389–3402. https://doi.org/10.1093/nar/25.17.3389.

Andrews, S., 2018. FastQC: a quality control tool for high throughput sequence data. https://www.bioinformatics.babraham.ac.uk/projects/fastqc/ [accessed 17 October 2022].

Arenas, M., Valiente, G., Posada, D., 2008. Characterization of reticulate networks based on the coalescent with recombination. Mol. Biol. Evol. 25, 2517–2520. https://doi.org/10.1093/molbev/msn219.

Bagley, J.C., Uribe-Convers, S., Carlsen, M.M., Muchhala, N., 2020. Utility of targeted sequence capture for phylogenomics in rapid, recent angiosperm radiations: Neotropical *Burmeistera* bellflowers as a case study. Mol. Phylogenet. Evol. 152, 106769. https://doi.org/10.1016/j.ympev.2020.106769.

Baker, W.J., Bailey, P., Barber, V., Barker, A., Bellot, S., Bishop, D., Botigué, L.R., Brewer, G., Carruthers, T., Clarkson, J.J., Cook, J., Cowan, R.S., Dodsworth, S., Epitawalage, N., Françoso, E., Gallego, B., Johnson, M.G., Kim, J.T., Leempoel, K., Maurin, O., Mcginnie, C., Pokorny, L., Roy, S., Stone, M., Toledo, E., Wickett, N.J., Zuntini, A.R., Eiserhardt, W.L., Kersey, P.J., Leitch, I.J., Forest, F., 2022. A comprehensive phylogenomic platform for exploring the angiosperm tree of life. Syst. Biol. 71, 301–319. https://doi.org/10.1093/sysbio/syab035.

Blischak, P.D., Chifman, J., Wolfe, A.D., Kubatko, L.S., 2018. HyDe: a Python package for genome-scale hybridization detection. Syst. Biol. 67(5), 821–829. https://doi.org/10.1093/sysbio/syy023.

Bolger, A.M., Lohse, M., Usadel, B., 2014. Trimmomatic: a flexible trimmer for Illumina sequence data. Bioinformatics 30, 2114–2120. https://doi.org/10.1093/bioinformatics/btu170.

Borowiec, M.L., 2016. AMAS: a fast tool for alignment manipulation and computing of summary statistics. PeerJ 4, e1660. https://doi.org/10.7717/peerj.1660.

Borowiec, M.L., 2019. Spruceup: fast and flexible identification, visualization, and removal of outliers from large multiple sequence alignments. J. Open Source Softw. 4, 1635. https://doi.org/10.21105/joss.01635.

Bouckaert, R., Heled, J., Kühnert, D., Vaughan, T., Wu, C.-H., Xie, D., Suchard, M.A., Rambaut, A., Drummond, A.J., 2014. BEAST 2: a software platform for Bayesian evolutionary analysis. PLoS Comput. Biol. 10, e1003537. https://doi.org/10.1371/journal.pcbi.1003537.

Brown, J.W., Walker, J.F., Smith, S.A., 2017. Phyx: phylogenetic tools for unix. Bioinformatics 33(12), 1886–1888. https://doi.org/10.1093/bioinformatics/btx063.

Byng, J.W., Chase, M.W., Christenhusz, M.J.M., Fay, M.F., Judd, W.S., Mabberley, D.J., Sennikov, A.N., Soltis, D.E., Soltis, P.S., Stevens, P.F., 2016. An update of the Angiosperm Phylogeny Group classification for the orders and families of flowering plants: APG IV. Bot. J. Linn. Soc. 181, 1–20. https://doi.org/10.1111/boj.12385.

Camacho, C., Coulouris, G., Avagyan, V., Ma, N., Papadopoulos, J., Bealer, K., Madden, T.L., 2009. BLAST+: architecture and applications. BMC Bioinformatics 10, 421. https://doi.org/10.1186/1471-2105-10-421.

Capella-Gutiérrez, S., Silla-Martínez, J.M., Gabaldón, T., 2009. trimAl: a tool for automated alignment trimming in large-scale phylogenetic analyses. Bioinformatics 25, 1972–1973. https://doi.org/10.1093/bioinformatics/btp348.

Chamala, S., García, N., Godden, G.T., Krishnakumar, V., Jordon-Thaden, I.E., De Smet, R., Barbazuk, W.B., Soltis, D.E., Soltis, P.S., 2015. MarkerMiner 1.0: a new application for phylogenetic marker development using angiosperm transcriptomes. Appl. Plant Sci. 3, apps.1400115. https://doi.org/10.3732/apps.1400115.

Christenhusz, M.J.M., Fay, M.F., Byng, J.W., 2018. The Global Flora A practical flora to vascular plant species of the world Special Edition, GLOVAP Nomenclature Part 1. Plant Gateway Ltd., Bradford, 4.

Cooper, B.J., Moore, M.J., Douglas, N.A., Wagner, W.L., Johnson, M.G., Overson, R.P., Kinosian, S.P., McDonnell, A.J., Levin, R.A., Raguso, R.A., Flores Olvera, H., Ochoterena, H., Fant, J.B., Skogen, K.A., Wickett, N.J., 2022. Target enrichment and extensive population sampling help untangle the recent, rapid radiation of *Oenothera* Sect. *Calylophus*. Syst. Biol. https://doi.org/10.1093/sysbio/syac032.

Crowl, A.A., Manos, P.S., McVay, J.D., Lemmon, A.R., Lemmon, E.M., Hipp, A.L., 2019. Uncovering the genomic signature of ancient introgression between white oak lineages (*Quercus*). New Phytol. 226, 1158–1170. https://doi.org/10.1111/nph.15842.

Dayrat, B., 2005. Towards integrative taxonomy. Biol. J. Linn. Soc. 85, 407–417. https://doi.org/10.1111/j.1095-8312.2005.00503.x.

Debray, K., Le Paslier, M.C., Berard, A., Thouroude, T., Michel, G., Marie-Magdelaine, J., Bruneau, A., Foucher, F., Malecot, V., 2022. Unveiling the patterns of reticulated evolutionary processes with phylogenomics: hybridization and polyploidy in the genus *Rosa*. Syst. Biol. 71, 547–569. https://doi.org/10.1093/sysbio/syab064.

Decaisne, M.J., 1874. Mémoire sur la famille des Pomacées. Nouvelles archives du muséum d’histoire naturelle 10, 113–192.

Dierckxsens, N., Mardulyn, P., Smits, G., 2016. Novoplasty: de novo assembly of organelle genomes from whole genome data. Nucleic Acids Res. 45, e18. https://doi.org/10.1093/nar/gkw955.

Flower, B.P., Kennett, J.P., 1994. The middle Miocene climatic transition: East Antarctic ice sheet development, deep ocean circulation and global carbon cycling. Paleogeogr. Paleoclimatol. Paleoecol. 108, 537–555. https://doi.org/10.1016/0031-0182(94)90251-8.

Focke, W.O., 1888. Rosaceae, in: Engler, A., Prantl, K. (Eds.), Die Naturlichen Pflanzenfamilien. Verlag von Wilhelm Engelmann, Leipzig, 3, 1–61.

Fonseca, L.H.M., Lohmann, L.G., 2020. Exploring the potential of nuclear and mitochondrial sequencing data generated through genome-skimming for plant phylogenetics: A case study from a clade of neotropical lianas. J. Syst. Evol. 58, 18–32. https://doi.org/10.1111/jse.12533.

Franchet, A.R., Delavay, J.M., 1890. Plantae Delavayanae. P. Klincksieck, Paris.

Guo, W., Yu, Y., Shen, R.J., Liao, W.B., Chin, S.-W., Potter, D., 2011. A phylogeny of *Photinia* sensu lato (Rosaceae) and related genera based on nrITS and cpDNA analysis. Plant Syst. Evol. 291(1), 91–102. https://doi.org/10.1007/s00606-010-0368-0.

Guo, W., Fan, Q., Zhang, X.Z., Liao, W.B., Wang, L.Y., Wu, W., Potter, D., 2020. Molecular reappraisal of relationships between *Photinia*, *Stranvaesia*, and *Heteromeles* (Rosaceae, Maleae). Phytotaxa 447, 103–115. https://doi.org/10.11646/phytotaxa.447.2.3.

Hodel, R.G.J., Zimmer, E.A., Liu, B.B., Wen, J., 2021. Synthesis of nuclear and chloroplast data combined with network analyses supports the polyploid origin of the apple tribe and the hybrid origin of the Maleae-Gillenieae clade. Front. Plant Sci. 12, 820997. https://doi.org/10.3389/fpls.2021.820997.

Hörandl, E., 2006. Paraphyletic versus monophyletic taxa-evolutionary versus cladistic classifications. Taxon 55, 564–570. https://doi.org/10.2307/25065631.

Huson, D.H., Bryant, D., 2006. Application of phylogenetic networks in evolutionary studies. Mol. Biol. Evol. 23, 254–267. https://doi.org/10.1093/molbev/msj030.

Huson, D.H., Scornavacca, C., 2011. A survey of combinatorial methods for phylogenetic networks. Genome Biol. Evol. 3, 23–35. https://doi.org/10.1093/gbe/evq077.

Jarvis, E.D., Mirarab, S., Aberer, A.J., Li, B., Houde, P., Li, C., Ho, S.Y.W., Faircloth, B.C., Nabholz, B., Howard, J.T., Suh, A., Weber, C.C., da Fonseca, R.R., Li, J.W., Zhang, F., Li, H., Zhou, L., Narula, N., Liu, L., Ganapathy, G., Boussau, B., Bayzid, M.S., Zavidovych, V., Subramanian, S., Gabaldón, T., Capella-Gutiérrez, S., Huerta-Cepas, J., Rekepalli, B., Munch, K., Schierup, M., Lindow, B., Warren, W.C., Ray, D., Green, R.E., Bruford, M.W., Zhan, X.J., Dixon, A., Li, S.B., Li, N., Huang, Y.H., Derryberry, E.P., Bertelsen, M.F., Sheldon, F.H., Brumfield, R.T., Mello, C.V., Lovell, P.V., Wirthlin, M., Schneider, M.P.C., Prosdocimi, F., Samaniego, J.A., Vargas Velazquez, A.M., Alfaro-Núñez, A., Campos, P.F., Petersen, B., Sicheritz-Ponten, T., Pas, A., Bailey, T., Scofield, P., Bunce, M., Lambert, D.M., Zhou, Q., Perelman, P., Driskell, A.C., Shapiro, B., Xiong, Z.J., Zeng, Y.L., Liu, S.P., Li, Z.Y., Liu, B.H., Wu, K., Xiao, J., Xiong, Y.Q., Zheng, Q.M., Zhang, Y., Yang, H.M., Wang, J., Smeds, L., Rheindt, F.E., Braun, M., Fjeldsa, J., Orlando, L., Barker, F.K., Jønsson, K.A., Johnson, W., Koepfli, K.-P., O’Brien, S., Haussler, D., Ryder, O.A., Rahbek, C., Willerslev, E., Graves, G.R., Glenn, T.C., McCormack, J., Burt, D., Ellegren, H., Alström, P., Edwards, S.V., Stamatakis, A., Mindell, D.P., Cracraft, J., Braun, E.L., Warnow, T., Jun, W., Gilbert, M.T.P., Zhang, G.J., 2014. Whole-genome analyses resolve early branches in the tree of life of modern birds. Science. 346, 1320–1331. https://doi.org/10.1126/science.1253451.

Johnson, M.G., Gardner, E.M., Liu, Y., Medina, R., Goffinet, B., Shaw, A.J., Zerega, N.J.C., Wickett, N.J., 2016. HybPiper: Extracting coding sequence and introns for phylogenetics from high-throughput sequencing reads using target enrichment. Appl. Plant Sci. 4, 1600016. https://doi.org/10.3732/apps.1600016.

Kalkman, C., 1973. The Malesian species of the subfamily Maloideae. Blumea 21, 413–442.

Kearse, M., Moir, R., Wilson, A., Stones-Havas, S., Cheung, M., Sturrock, S., Buxton, S., Cooper, A., Markowitz, S., Duran, C., Thierer, T., Ashton, B., Meintjes, P., Drummond, A., 2012. Geneious Basic: an integrated and extendable desktop software platform for the organization and analysis of sequence data. Bioinformatics 28, 1647–1649. https://doi.org/10.1093/bioinformatics/bts199.

Koehne, E., 1890. Die Gattungen der Pomaceen. Wissenschaftliche Beilage zum Programm des Falk-Realgymnasiums zu, Berlin, pp. 1–33.

Koehne, E., 1891. Die Gattungen der Pomaceen, in: Engler, A., Prantl, K. (Eds.), Garten flora. Verlag von Paul Parey, Berlin, 40, 4–7.

Lanfear, R., Calcott, B., Ho, S.Y., Guindon, S., 2012. PartitionFinder: combined selection of partitioning schemes and substitution models for phylogenetic analyses. Mol. Biol. Evol. 29(6), 1695–1701. https://doi.org/10.1093/molbev/mss020.

Lanfear, R., Calcott, B., Kainer, D., Mayer, C., Stamatakis, A., 2014. Selecting optimal partitioning schemes for phylogenomic datasets. BMC Evol. Biol. 14, 82. https://doi.org/10.1186/1471-2148-14-82.

Lanfear, R., Frandsen, P.B., Wright, A.M., Senfeld, T., Calcott, B., 2016. PartitionFinder 2: new methods for selecting partitioned models of evolution for molecular and morphological phylogenetic analyses. Mol. Biol. Evol. 34, 772–773. https://doi.org/10.1093/molbev/msw260.

Larget, B.R., Kotha, S.K., Dewey, C.N., Ané, C., 2010. BUCKy: gene tree/species tree reconciliation with Bayesian concordance analysis. Bioinformatics 26(22), 2910–2911. https://doi.org/10.1093/bioinformatics/btq539.

Leebens-Mack, J.H., Barker, M.S., Carpenter, E.J., Deyholos, M.K., Gitzendanner, M.A., Graham, S.W., Grosse, I., Li, Z., Melkonian, M., Mirarab, S., Porsch, M., Quint, M., Rensing, S.A., Soltis, D.E., Soltis, P.S., Stevenson, D.W., Ullrich, K.K., Wickett, N.J., DeGironimo, L., Edger, P.P., Jordon-Thaden, I.E., Joya, S., Liu, T., Melkonian, B., Miles, N.W., Pokorny, L., Quigley, C., Thomas, P., Villarreal, J.C., Augustin, M.M., Barrett, M.D., Baucom, R.S., Beerling, D.J., Benstein, R.M., Biffin, E., Brockington, S.F., Burge, D.O., Burris, J.N., Burris, K.P., Burtet-Sarramegna, V., Caicedo, A.L., Cannon, S.B., Çebi, Z., Chang, Y., Chater, C., Cheeseman, J.M., Chen, T., Clarke, N.D., Clayton, H., Covshoff, S., Crandall-Stotler, B.J., Cross, H., dePamphilis, C.W., Der, J.P., Determann, R., Dickson, R.C., Di Stilio, V.S., Ellis, S., Fast, E., Feja, N., Field, K.J., Filatov, D.A., Finnegan, P.M., Floyd, S.K., Fogliani, B., García, N., Gâteblé, G., Godden, G.T., Goh, F., Greiner, S., Harkess, A., Heaney, J.M., Helliwell, K.E., Heyduk, K., Hibberd, J.M., Hodel, R.G.J., Hollingsworth, P.M., Johnson, M.T.J., Jost, R., Joyce, B., Kapralov, M.V., Kazamia, E., Kellogg, E.A., Koch, M.A., Von Konrat, M., Könyves, K., Kutchan, T.M., Lam, V., Larsson, A., Leitch, A.R., Lentz, R., Li, F.-W., Lowe, A.J., Ludwig, M., Manos, P.S., Mavrodiev, E., McCormick, M.K., McKain, M., McLellan, T., McNeal, J.R., Miller, R.E., Nelson, M.N., Peng, Y., Ralph, P., Real, D., Riggins, C.W., Ruhsam, M., Sage, R.F., Sakai, A.K., Scascitella, M., Schilling, E.E., Schlösser, E.-M., Sederoff, H., Servick, S., Sessa, E.B., Shaw, A.J., Shaw, S.W., Sigel, E.M., Skema, C., Smith, A.G., Smithson, A., Stewart, C.N., Stinchcombe, J.R., Szövényi, P., Tate, J.A., Tiebel, H., Trapnell, D., Villegente, M., Wang, C.-N., Weller, S.G., Wenzel, M., Weststrand, S., Westwood, J.H., Whigham, D.F., Wu, S.X., Wulff, A.S., Yang, Y., Zhu, D., Zhuang, C.L., Zuidof, J., Chase, M.W., Pires, J.C., Rothfels, C.J., Yu, J., Chen, C., Chen, L., Cheng, S.F., Li, J.J., Li, R., Li, X., Lu, H.R., Ou, Y.X., Sun, X., Tan, X.M., Tang, J.B., Tian, Z.J., Wang, F., Wang, J., Wei, X.F., Xu, X., Yan, Z.X., Yang, F., Zhong, X.N., Zhou, F.Y., Zhu, Y., Zhang, Y., Ayyampalayam, S., Barkman, T.J., Nguyen, N.-p., Matasci, N., Nelson, D.R., Sayyari, E., Wafula, E.K., Walls, R.L., Warnow, T., An, H., Arrigo, N., Baniaga, A.E., Galuska, S., Jorgensen, S.A., Kidder, T.I., Kong, H.H., Lu-Irving, P., Marx, H.E., Qi, X.S., Reardon, C.R., Sutherland, B.L., Tiley, G.P., Welles, S.R., Yu, R.P., Zhan, S., Gramzow, L., Theißen, G., Wong, G.K.-S., 2019. One thousand plant transcriptomes and the phylogenomics of green plants. Nature 574, 679–685. https://doi.org/10.1038/s41586-019-1693-2.

Lehwark, P., Greiner, S., 2019. GB2sequin - A file converter preparing custom GenBank files for database submission. Genomics 111(4), 759–761. https://doi.org/10.1016/j.ygeno.2018.05.003.

Li, G., Lu, L.T., Li, C.L., 1992. Leaf architecture of the *Photinia* complex (Rosaceae: Maloideae) with special reference to its phenetic and phylogenetic significance. Cathaya 3, 21–56.

Li, H.T., Yi, T.S., Gao, L.M., Ma, P.F., Zhang, T., Yang, J.B., Gitzendanner, M.A., Fritsch, P.W., Cai, J., Luo, Y., Wang, H., van der Bank, M., Zhang, S.D., Wang, Q.F., Wang, J., Zhang, Z.R., Fu, C.N., Yang, J., Hollingsworth, P.M., Chase, M.W., Soltis, D.E., Soltis, P.S., Li, D.Z., 2019. Origin of angiosperms and the puzzle of the Jurassic gap. Nat. Plants 5, 461. https://doi.org/10.1038/s41477-019-0421-0.

Li, H.T., Luo, Y., Gan, L., Ma, P.F., Gao, L.M., Yang, J.B., Cai, J., Gitzendanner, M.A., Fritsch, P.W., Zhang, T., Jin, J.J., Zeng, C.X., Wang, H., Yu, W.B., Zhang, R., van der Bank, M., Olmstead, R.G., Hollingsworth, P.M., Chase, M.W., Soltis, D.E., Soltis, P.S., Yi, T.S., Li, D.Z., 2021. Plastid phylogenomic insights into relationships of all flowering plant families. BMC Biol. 19, 232. https://doi.org/10.1186/s12915-021-01166-2.

Li, J.L., Wang, S., Yu, J., Wang, L., Zhou, S.L., 2013. A modified CTAB protocol for plant DNA extraction. Chin. Bull. Bot. 48, 72–78 (in Chinese).

Liang, G.L., 1986. Comparative studies of karyotypes in Chinese species of *Malus*. J. Southwest Agric. Univ. 1, 104–117 (in Chinese).

Liang, G.L., 1987. Observations of chromosomes of *Malus* species in China. Acta Phytotax. Sin. 25, 437–441 (in Chinese).

Liang, G.L., Li, X.L., 1993. Chromosome studies of Chinese species of *Malus* Mill. Acta Phytotax. Sin. 31, 236–251 (in Chinese).

Liang, G.L., Li, Y.N., Li, X.L., 1996. Evolutionary study of the chromosomes at pollen mother cell meiosis in *Malus*. J. Southwest Agric. Univ. 18, 299–307 (in Chinese).

Liang, G.L., 1997. Induction of Giemsa C-bands and analyses of banding patterns in *Malus sikkimensis*. J. Southwest Agric. Univ. 19, 95–97 (in Chinese).

Liu, B.B., Campbell, C.S., Hong, D.Y., Wen, J., 2020a. Phylogenetic relationships and chloroplast capture in the *Amelanchier*-*Malacomeles*-*Peraphyllum* clade (Maleae, Rosaceae): Evidence from chloroplast genome and nuclear ribosomal DNA data using genome skimming. Mol. Phylogenet. Evol. 147, 106784. https://doi.org/10.1016/j.ympev.2020.106784.

Liu, B.B., Hong, D.Y., Zhou, S.L., Xu, C., Dong, W.P., Johnson, G., Wen, J., 2019. Phylogenomic analyses of the *Photinia* complex support the recognition of a new genus *Phippsiomeles* and the resurrection of a redefined *Stranvaesia* in Maleae (Rosaceae). J. Syst. Evol. 57, 678–694. https://doi.org/10.1111/jse.12542.

Liu, B.B., Liu, G.N., Hong, D.Y., Wen, J., 2020b. *Eriobotrya* belongs to *Rhaphiolepis* (Maleae, Rosaceae): evidence from chloroplast genome and nuclear ribosomal DNA data. Front. Plant Sci. 10, 1731. https://doi.org/10.3389/fpls.2019.01731.

Liu, B.B., Ma, Z.Y., Ren, C., Hodel, R.G.J., Sun, M., Liu, X.Q., Liu, G.N., Hong, D.Y., Zimmer, E.A., Wen, J., 2021. Capturing single-copy nuclear genes, organellar genomes, and nuclear ribosomal DNA from deep genome skimming data for plant phylogenetics: A case study in Vitaceae. J. Syst. Evol. 59, 1124–1138. https://doi.org/10.1111/jse.12806.

Liu, B.B., Ren, C., Kwak, M., Hodel, R.G.J., Xu, C., He, J., Zhou, W.B., Huang, C.-H., Ma, H., Qian, G.Z., Hong, D.Y., Wen, J., 2022. Phylogenomic conflict analyses in the apple genus *Malus* s.l. reveal widespread hybridization and allopolyploidy driving diversification, with insights into the complex biogeographic history in the Northern Hemisphere. J. Integr. Plant Biol. 64(5), 1020– 1043. https://doi.org/10.1111/jipb.13246.

Liu, X., Wang, Z.S., Shao, W.H., Ye, Z.Y., Zhang, J.G., 2017. Phylogenetic and taxonomic status analyses of the Abaso Section from multiple nuclear genes and plastid fragments reveal new insights into the North America origin of *Populus* (Salicaceae). Front. Plant Sci. 7, 2022. https://doi.org/10.3389/fpls.2016.02022.

Lo, E.Y.Y., Donoghue, M.J., 2012. Expanded phylogenetic and dating analyses of the apples and their relatives (Pyreae, Rosaceae). Mol. Phylogenet. Evol. 63, 230–243. https://doi.org/10.1016/j.ympev.2011.10.005.

Lu, L.T., Li, C.L., Li, G., 1990. Pollen morphology of *Photinia* (Rosaceae)and its systematic significance. Cathaya 2, 127–138.

Lu, L.T., Wang, Z.L., Li, G., 1991. The significance of the leaf epidermis in the taxonomy of the *Photinia* complex (Rosaceae: Maloideae). Cathaya 3, 93–108.

Lu, L.T., Spongberg, S.A., 2003. *Photinia* Lindley, in: Wu, Z.Y., Raven, P.H., Hong, D.Y. (Eds.), Flora of China. Vol. 9. Science Press, Beijing; Missouri Botanical Garden Press, St. Louis, pp. 121–137.

MacGinitie, H.D., 1969. The Eocene Green River Flora of northwestern Colorado and northeastern Utah. Univ. Calif. Publ. Geol. Sci. 83, 1–140.

Mai, U., Mirarab, S., 2018. TreeShrink: fast and accurate detection of outlier long branches in collections of phylogenetic trees. BMC Genomics 19(S5), 272. https://doi.org/10.1186/s12864-018-4620-2.

Mallet, J., 2005. Hybridization as an invasion of the genome. Trends Ecol. Evol. 20, 229–237. https://doi.org/10.1016/j.tree.2005.02.010.

Matzke, N.J., 2018. BioGeoBEARS: BioGeography with Bayesian (and likelihood) Evolutionary Analysis with R Scripts. version 1.1.1, published on GitHub on November 6, 2018. http://dx.doi.org/10.5281/zenodo.1478250.

Mayr, E., Bock, W.J., 2002. Classifications and other ordering systems. J. Zool. Syst. Evol. Res. 40, 169–194. https://doi.org/10.1046/j.1439-0469.2002.00211.x.

Mayrose, I., Zhan, S.H., Rothfels, C.J., Magnuson-Ford, K., Barker, M.S., Rieseberg, L.H., Otto, S.P., 2011. Recently formed polyploid plants diversify at lower rates. Science 333, 1257. https://doi.org/10.1126/science.1207205.

Mehra, P.N., Sareen, T.S., Hans, A.S., 1973. Cytology of some woody species of Rosaceae from Himalayas. Silvae Genet. 22, 188–190.

Mindell, D.P., 2013. The tree of life: metaphor, model, and heuristic device. Syst. Biol. 62, 479–489. https://doi.org/10.1093/sysbio/sys115.

Minh, B.Q., Schmidt, H.A., Chernomor, O., Schrempf, D., Woodhams, M.D., Von Haeseler, A., Lanfear, R., 2020. IQ-TREE 2: New models and efficient methods for phylogenetic inference in the genomic era. Mol. Biol. Evol. 37, 1530–1534. https://doi.org/10.1093/molbev/msaa015.

Morales-Briones, D.F., Kadereit, G., Tefarikis, D.T., Moore, M.J., Smith, S.A., Brockington, S.F., Timoneda, A., Yim, W.C., Cushman, J.C., Yang, Y., 2021. Disentangling sources of gene tree discordance in phylogenomic data sets: testing ancient hybridizations in Amaranthaceae s.l. Syst. Biol. 70, 219–235. https://doi.org/10.1093/sysbio/syaa066.

Morales-Briones, D.F., Gehrke, B., Huang, C.-H., Liston, A., Ma, H., Marx, H.E., Tank, D.C., Yang, Y., 2022. Analysis of paralogs in target enrichment data pinpoints multiple ancient polyploidy events in *Alchemilla* s.l. (Rosaceae). Syst. Biol. 71, 190–207. https://doi.org/10.1093/sysbio/syab032.

Morgulis, A., Coulouris, G., Raytselis, Y., Madden, T.L., Agarwala, R., Schäffer, A.A., 2008. Database indexing for production Megablast searches. Bioinformatics 24, 1757–1764. https://doi.org/10.1093/bioinformatics/btn322.

Morrison, D.A., 2014. Is the tree of life the best metaphor, model, or heuristic for phylogenetics? Syst. Biol. 63, 628–638. https://doi.org/10.1093/sysbio/syu026.

Nakamura, T., Yamada, K.D., Tomii, K., Katoh, K., 2018. Parallelization of MAFFT for large-scale multiple sequence alignments. Bioinformatics 34, 2490–2492. https://doi.org/10.1093/bioinformatics/bty121.

Padial, J.M., Miralles, A., De la Riva, I., Vences, M., 2010. The integrative future of taxonomy. Front. Zool. 7, 1–14. https://doi.org/10.1186/1742-9994-7-16.

Padian, K., 1999. Charles Darwin’s views of classification in theory and practice. Syst. Biol. 48, 352–364. https://doi.org/10.1080/106351599260337.

Pathak, M.L., Idrees, M., Xu, B., Gao, X.F., 2019. Pollen morphology of subfamily Maloideae (Roseaceae) with special focus on the genus *Photinia*. Phytotaxa 404(5), 171–189. https://doi.org/10.11646/phytotaxa.404.5.1.

Pease, J.B., Brown, J.W., Walker, J.F., Hinchliff, C.E., Smith, S.A., 2018. Quartet sampling distinguishes lack of support from conflicting support in the green plant tree of life. Am. J. Bot. 105, 385–403. https://doi.org/10.1002/ajb2.1016.

Rehder, A., 1940. Manual of cultivated trees and shrubs hardy in North America exclusive of the subtropical and warmer temperature regions, 2nd ed. The Macmillan Company, New York.

Rehder, A., 1949. Bibliography of cultivated trees and shrubs hardy in the cooler temperature regions of the Northern Hemisphere. The Arnold Arboretum of Harvard University, Jamaica Plain, Massachusetts.

Rieseberg, L.H., Soltis, D.E., 1991. Phylogenetic consequences of cytoplasmic gene flow in plants. Evol. Trends Plants 5, 65–84.

Rieseberg, L.H., Willis, J.H., 2007. Plant speciation. Science. 317, 910–914. https://doi.org/10.1126/science.1137729.

Robertson, K.R., Phipps, J.B., Rohrer, J.R., Smith, P.G., 1991. A synopsis of genera in Maloideae (Rosaceae). Syst. Bot. 16, 376–394. https://doi.org/10.2307/2419287.

Roemer, M.J., 1847. Familiarum naturalium regni vegetabilis synopses monographicae. Vol. 3. Landes-Industrie-Comptoir, Weimar.

Rose, J.P., Toledo, C.A.P., Lemmon, E.M., Lemmon, A.R., Sytsma, K.J., 2020. Out of sight, out of mind: widespread nuclear and plastid-nuclear discordance in the flowering plant genus *Polemonium* (Polemoniaceae) suggests widespread historical gene flow despite limited nuclear signal. Syst. Biol. 70, 162–180. https://doi.org/10.1093/sysbio/syaa049.

Rothfels, C.J., 2021. Polyploid phylogenetics. New Phytol. 230, 66–72. https://doi.org/10.1111/nph.17105.

Sayyari, E., Mirarab, S., 2016. Fast coalescent-based computation of local branch support from quartet frequencies. Mol. Biol. Evol. 33(7), 1654–1668. https://doi.org/10.1093/molbev/msw079.

Schlick-Steiner, B.C., Steiner, F.M., Seifert, B., Stauffer, C., Christian, E., Crozier, R.H., 2010. Integrative taxonomy: a multisource approach to exploring biodiversity. Annu. Rev. Entomol. 55, 421–438. https://doi.org/10.1146/annurev-ento-112408-085432.

Schliep, K., Potts, A.J., Morrison, D.A., Grimm, G.W., 2017. Intertwining phylogenetic trees and networks. Methods Ecol. Evol. 8, 1212–1220. https://doi.org/10.1111/2041-210X.12760.

Schuettpelz, E., Schneider, H., Smith, A.R., Hovenkamp, P., Prado, J., Rouhan, G., Salino, A., Sundue, M., Almeida, T.E., Parris, B., Sessa, E.B., Field, A.R., de Gasper, A.L., Rothfels, C.J., Windham, M.D., Lehnert, M., Dauphin, B., Ebihara, A., Lehtonen, S., Schwartsburd, P.B., Metzgar, J., Zhang, L.B., Kuo, L.-Y., Brownsey, P.J., Kato, M., Arana, M.D., 2016. A community-derived classification for extant lycophytes and ferns. J. Syst. Evol. 54, 563–603. https://doi.org/10.1111/jse.12229.

Singhal, V.K., Gill, B.S., Sidhu, M.S., 1990. Cytology of woody members of Rosaceae. Proc. Plant Sci. 100, 17–21. https://doi.org/10.1007/BF03053464.

Smith, S.A., O’Meara, B.C., 2012. treePL: divergence time estimation using penalized likelihood for large phylogenies. Bioinformatics 28(20), 2689–2690. https://doi.org/10.1093/bioinformatics/bts492.

Smith, S.A., Moore, M.J., Brown, J.W., Yang, Y., 2015. Analysis of phylogenomic datasets reveals conflict, concordance, and gene duplications with examples from animals and plants. BMC Evol. Biol. 15, 150. https://doi.org/10.1186/s12862-015-0423-0.

Smith, B.T., Merwin, J., Provost, K.L., Thom, G., Brumfield, R.T., Ferreira, M., Mauck Iii, W.M., Moyle, R.G., Wright, T.F., Joseph, L., 2022. Phylogenomic analysis of the parrots of the world distinguishes artifactual from biological sources of gene tree discordance. Syst. Biol. https://dx.doi.org/10.1093/sysbio/syac055.

Solís-Lemus, C., Ané, C., 2016. Inferring phylogenetic networks with maximum pseudolikelihood under incomplete lineage sorting. PLoS Genet. 12(3), e1005896. https://doi.org/10.1371/journal.pgen.1005896.

Solís-Lemus, C., Bastide, P., Ané, C., 2017. PhyloNetworks: A package for phylogenetic networks. Mol. Biol. Evol. 34, 3292–3298. https://doi.org/10.1093/molbev/msx235.

Soltis, D.E., Soltis, P.S., Collier, T.G., Edgerton, M.L., 1991. Chloroplast DNA variation within and among genera of the *Heuchera* group (Saxifragaceae): evidence for chloroplast transfer and paraphyly. Am. J. Bot. 78, 1091–1112. https://doi.org/10.1002/j.1537-2197.1991.tb14517.x.

Stamatakis, A., 2006. RAxML-VI-HPC: maximum likelihood-based phylogenetic analyses with thousands of taxa and mixed models. Bioinformatics 22, 2688–2690. https://doi.org/10.1093/bioinformatics/btl446.

Stamatakis, A., 2014. RAxML version 8: a tool for phylogenetic analysis and post-analysis of large phylogenies. Bioinformatics 30, 1312–1313. https://doi.org/10.1093/bioinformatics/btu033.

Straub, S.C.K., Parks, M., Weitemier, K., Fishbein, M., Cronn, R.C., Liston, A., 2012. Navigating the tip of the genomic iceberg: Next-generation sequencing for plant systematics. Am. J. Bot. 99, 349–364. https://doi.org/10.3732/ajb.1100335.

Su, N., Liu, B.B., Wang, J.R., Tong, R.C., Ren, C., Chang, Z.Y., Zhao, L., Potter, D., Wen, J., 2021. On the species delimitation of the *Maddenia* group of *Prunus* (Rosaceae): evidence from plastome and nuclear sequences and morphology. Front. Plant Sci. 12, 743643. https://doi.org/10.3389/fpls.2021.743643.

Thomas, G.W.C., Ather, S.H., Hahn, M.W., 2017. Gene-tree reconciliation with MUL-trees to resolve polyploid events. Syst. Biol. 66(6), 1007–1018. https://doi.org/10.1093/sysbio/syx044.

Upham, N.S., Esselstyn, J.A., Jetz, W., 2019. Inferring the mammal tree: Species-level sets of phylogenies for questions in ecology, evolution, and conservation. PLoS Biol. 17, e3000494. https://doi.org/10.1371/journal.pbio.3000494.

Weitemier, K., Straub, S.C.K., Fishbein, M., Liston, A., 2015. Intragenomic polymorphisms among high-copy loci: a genus-wide study of nuclear ribosomal DNA in *Asclepias* (Apocynaceae). PeerJ 3, e718. https://doi.org/10.7717/peerj.718.

Wen, J., Harris, A.J., Ickert-Bond, S.M., Dikow, R., Wurdack, K., Zimmer, E.A., 2017. Developing integrative systematics in the informatics and genomic era, and calling for a global Biodiversity Cyberbank. J. Syst. Evol. 55, 308–321. https://doi.org/10.1111/jse.12270.

Wen, J., Xiong, Z.Q., Nie, Z.L., Mao, L.K., Zhu, Y.B., Kan, X.Z., Ickert-Bond, S.M., Gerrath, J., Zimmer, E.A., Fang, X.D., 2013. Transcriptome sequences resolve deep relationships of the grape family. PLoS One 8, e74394. https://doi.org/10.1371/journal.pone.0074394.

Wen, D.Q., Yu, Y., Zhu, J.F., Nakhleh, L., 2018. Inferring phylogenetic networks using PhyloNet. Syst. Biol. 67, 735–740. https://doi.org/10.1093/sysbio/syy015.

Wenzig, T., 1883. Die Pomaceen, Charaktere der Gattungen und Arten. Vol. 2. Jahrbuch des Königlichen Botanischen Gartens und des Botanischen Museums zu Berlin, Berlin, pp. 287–307.

Wickett, N.J., Mirarab, S., Nguyen, N., Warnow, T., Carpenter, E., Matasci, N., Ayyampalayam, S., Barker, M.S., Burleigh, J.G., Gitzendanner, M.A., Ruhfel, B.R., Wafula, E., Der, J.P., Graham, S.W., Mathews, S., Melkonian, M., Soltis, D.E., Soltis, P.S., Miles, N.W., Rothfels, C.J., Pokorny, L., Shaw, A.J., DeGironimo, L., Stevenson, D.W., Surek, B., Villarreal, J.C., Roure, B., Philippe, H., dePamphilis, C.W., Chen, T., Deyholos, M.K., Baucom, R.S., Kutchan, T.M., Augustin, M.M., Wang, J., Zhang, Y., Tian, Z.J., Yan, Z.X., Wu, X.L., Sun, X., Wong, G.K.-S., Leebens-Mack, J., 2014. Phylotranscriptomic analysis of the origin and early diversification of land plants. Proc. Natl. Acad. Sci. USA 111, E4859–E4868. https://doi.org/10.1073/pnas.1323926111.

Wood, T.E., Takebayashi, N., Barker, M.S., Mayrose, I., Greenspoon, P.B., Rieseberg, L.H., 2009. The frequency of polyploid speciation in vascular plants. Proc. Natl. Acad. Sci. USA 106(33), 13875– 13879. https://doi.org/10.1073/pnas.0811575106.

Xiang, Y.Z., Huang, C.-H., Hu, Y., Wen, J., Li, S.S., Yi, T.S., Chen, H.Y., Xiang, J., Ma, H., 2017. Evolution of Rosaceae fruit types based on nuclear phylogeny in the context of geological times and genome duplication. Mol. Biol. Evol. 34, 262–281. https://doi.org/10.1093/molbev/msw242.

Yang, Y., Smith, S.A., 2014. Orthology inference in nonmodel organisms using transcriptomes and low-coverage genomes: improving accuracy and matrix occupancy for phylogenomics. Mol. Biol. Evol. 31, 3081–3092. https://doi.org/10.1093/molbev/msu245.

Yang, Y.Z., Sun, P.C., Lv, L.K., Wang, D.L., Ru, D.F., Li, Y., Ma, T., Zhang, L., Shen, X.X., Meng, F.B., Jiao, B.B., Shan, L.X., Liu, M., Wang, Q.F., Qin, Z.J., Xi, Z.X., Wang, X.Y., Davis, C.C., Liu, J.Q., 2020. Prickly waterlily and rigid hornwort genomes shed light on early angiosperm evolution. Nat. Plants 6, 215–222. https://doi.org/10.1038/s41477-020-0594-6.

Yi, T.S., Jin, G.H., Wen, J., 2015. Chloroplast capture and intra- and inter-continental biogeographic diversification in the Asian-New World disjunct plant genus *Osmorhiza* (Apiaceae). Mol. Phylogenet. Evol. 85, 10–21. https://doi.org/10.1016/j.ympev.2014.09.028.

Yu, T.T., Ku, T.C., 1974. *Stranvaesia* Lindl, in: Yu, T.T. (Ed.), Flora Reipublicae Popularis Sinicae. Vol. 36. Science Press, Beijing, pp. 210–216.

Yu, Y., Harris, A.J., Blair, C., He, X.J., 2015. RASP (Reconstruct Ancestral State in Phylogenies): a tool for historical biogeography. Mol. Phylogenet. Evol. 87, 46–49. https://doi.org/10.1016/j.ympev.2015.03.008.

Zhang, C., Rabiee, M., Sayyari, E., Mirarab, S., 2018. ASTRAL-III: polynomial time species tree reconstruction from partially resolved gene trees. BMC Bioinformatics 19, 153. https://doi.org/10.1186/s12859-018-2129-y.

Zhang, N., Wen, J., Zimmer, E.A., 2015. Congruent deep relationships in the grape family (Vitaceae) based on sequences of chloroplast genomes and mitochondrial genes via genome skimming. PLoS One 10, e0144701. https://doi.org/10.1371/journal.pone.0144701.

Zhang, S.Y., 1992. Systematic wood anatomy of the Rosaceae. Blumea: Biodiversity, Evolution and Biogeography of Plants 37, 81–158.

Zhang, S.D., Jin, J.J., Chen, S.Y., Chase, M.W., Soltis, D.E., Li, H.T., Yang, J.B., Li, D.Z., Yi, T.S., 2017. Diversification of Rosaceae since the Late Cretaceous based on plastid phylogenomics. New Phytol. 214(3), 1355–1367. https://doi.org/10.1111/nph.14461.

Zhao, M., Kurtis, S.M., White, N.D., Moncrieff, A.E., Leite, R.N., Brumfield, R.T., Braun, E.L., Kimball, R.T., 2022. Exploring conflicts in Whole Genome Phylogenetics: A case study within Manakins (Aves: Pipridae). Syst. Biol. https://doi.org/10.1093/sysbio/syac062.

Zhou, B.F., Yuan, S., Crowl, A.A., Liang, Y.Y., Shi, Y., Chen, X.Y., An, Q.Q., Kang, M., Manos, P.S., Wang, B.S. 2022. Phylogenomic analyses highlight innovation and introgression in the continental radiations of Fagaceae across the Northern Hemisphere. Nat. Commun. 13, 1320. https://doi.org/10.1038/s41467-022-28917-1.

Zimmer, E.A., Wen, J., 2015. Using nuclear gene data for plant phylogenetics: Progress and prospects II. Next-gen approaches. J. Syst. Evol. 53, 371–379. https://doi.org/10.1111/jse.12174.

Zou, X.H., Ge, S., 2008. Conflicting gene trees and phylogenomics. J. Syst. Evol. 46(6), 795. https://doi.org/10.1093/sysbio/syz078.

