## Supplementary_material for "Nightmare or delight: taxonomic circumscription meets reticulate evolution in the phylogenomic era": Appendix A. Supplementary material caption.pdf

Supplementary Fig. S1. Heat map showing percentage length recovery for 801 genes recovered by HybPiper. Each row shows a sample, and each column is a gene. The amount of shading in each box corresponds to the length of the gene recovered for that sample by the pipeline, relative to mean reference length (maximum of 1.0).

Supplementary Fig. S2. Maximum likelihood phylogeny of *Stranvaesia* within Maleae inferred from RAxML analysis of plastid CDS dataset. Bootstrap support (BS) is shown above branches.

Supplementary Fig. S3. Maximum likelihood phylogeny of *Stranvaesia* within Maleae inferred from IQ-TREE2 analysis of plastid CDS dataset. the SH-aLRT support and Ultrafast Bootstrap support are shown above branches.

Supplementary Fig. S4. ASTRAL-III Species tree of *Stranvaesia* within Maleae inferred from CDS dataset. Local posterior probabilities (LLP) are shown above branches.

Supplementary Fig. S5. Maximum likelihood phylogeny of *Stranvaesia* within Maleae inferred from RAxML analysis of CDS dataset. The number of gene trees concordant/conflicting with that node in the nuclear phylogeny are shown above branches. The Internode Certainty All (ICA) score are shown below branches. Pie charts on nodes denote the proportion of gene trees that support that clade (blue), the proportion that support the main alternative bifurcation (green), the proportion that support the remaining alternatives (red), and the proportion (conflict or support) that have < 50% bootstrap support (gray).

Supplementary Fig. S6. Maximum likelihood phylogeny of *Stranvaesia* within Maleae inferred from RAxML analysis of CDS dataset. Quartet Sampling (QS) scores for each node are shown next to branches indicating Quartet Concordance (QC) / Quartet Differential (QD) / Quartet Informativeness (QI). Quartet Concordance is also showed in each node'pie chart and color-coded according to the legend.

Supplementary Fig. S7. Maximum likelihood phylogeny of *Stranvaesia* within Maleae inferred from RAxML analysis of SCNs dataset. Bootstrap support (BS) is shown above branches

Supplementary Fig. S8. Maximum likelihood phylogeny of *Stranvaesia* within Maleae inferred from IQ-TREE2 analysis of SCNs dataset. the SH-aLRT support and Ultrafast Bootstrap support are shown above branches.

Supplementary Fig. S9. ASTRAL-III Species tree of *Stranvaesia* within Maleae inferred from SCNs dataset. Local posterior probabilities (LLP) are shown above branches.

Supplementary Fig. S10. Maximum likelihood phylogeny of *Stranvaesia* within Maleae inferred from RAxML analysis of SCNs dataset. The number of gene trees concordant/conflicting with that node in the nuclear phylogeny are shown above branches. The Internode Certainty All (ICA) score are shown below branches. Pie charts on nodes denote the proportion of gene trees that support that clade (blue), the proportion that support the main alternative bifurcation (green), the proportion that support the remaining alternatives (red), and the proportion (conflict or support) that have < 50% bootstrap support (gray).

Supplementary Fig. S11. Maximum likelihood phylogeny of *Stranvaesia* within Maleae inferred from RAxML analysis of SCNs dataset. Quartet Sampling (QS) scores for each node are shown next to branches indicating Quartet Concordance (QC) / Quartet Differential (QD) / Quartet Informativeness (QI). Quartet Concordance is also showed in each node'pie chart and color-coded according to the legend.

Supplementary Fig. S12. Phylogenetic network analysis from the 12-taxa sampling of *Stranvaesia* and its closely relative genera. Species networks inferred from SNaQ network analysis allowing for zero ($h_{max}$ = 0) to five ($h_{max}$ = 5) hybridization events, except for $h_{max}$ = 2, which was considered the best network. Blue curved branches indicate the possible hybridization event. Dark blue and light blue numbers indicate the major and minor inheritance probabilities of hybrid nodes.

Supplementary Fig. S13. The most parsimonious multi-labeled trees inferred from GRAMPA analyses on the trees inferred from nuclear phylogeny with the *Stranvaesia* clade set as a result of allopolyploidization. The clade with multiple labels denotes the polyploidy origin, a plus sign indicates the first tip, and the second tip is shown with an asterisk. One of the *Stranvaesia* clade is sister to the *Weniomeles* clade, and the other of the *Stranvaesia* clade is sister to the ancestor of *Stranvaesia*, indicating the allopolyploid origin of the *Stranvaesia* clade.

Supplementary Fig. S14. Dated chronogram for the red-fruit genus *Stranvaesia* inferred from treePL based on the 426 SCN genes dataset.

Supplementary Fig. S15. The ancestral area reconstruction using BioGeoBEARS implemented in RASP from the 426 SCN genes dataset with fossil record only used for calibration in the framework of Maleae, with the colored key identifying extant and possible ancestral ranges. (A), East Asia; (B), Europe; (C), Central Asia; (D), North America; (E), South America.

Supplementary Table S1. Voucher and sequences information for taxa of the *Stranvaesia* and its relatives used in this study.

Supplementary Table S2. The number of pre-filtered and post-filtered sequences for each sample in this study.
