## Supplementary_material for "Nightmare or delight: taxonomic circumscription meets reticulate evolution in the phylogenomic era": SUPPLEMENTARY.TABLE.S1.PDF.pdf

Table S1. Voucher and sequences information for taxa of the *Stranvaesia* and its relatives used in this study

| Species | Type of data | No. of clean reads | Sequencing depth | Locality | Voucher | NCBI accession | Plastome accession |
|---|---|---|---|---|---|---|---|
| *Amelanchier alnifolia* | RNA-Seq | 122,179,876 | 24.4 | Pennsylvania, USA | HM976 (FUS) | SRR15691199 | MN068255 |
| *Aronia melanocarpa* | RNA-Seq | 40,313,330 | 8.1 | USA | - | SRR10963617 | KY420007 |
| *Chaenomeles japonica* | DGS | 46,818,834 | 9.4 | Beijing, China | B.B.Liu & G.N.Liu 3926 (PE) | SRR15691198 | MZ984211 |
| *Chaenomeles speciosa* | DGS | 52,357,252 | 10.5 | Beijing, China | B.B.Liu 3985 (PE) | SRR15691187 | MZ984212 |
| *Cormus domestica* | DGS | 46,534,300 | 9.3 | USA | B.B.Liu Z0925 | SRR15691176 | MZ984208 |
| *Cotoneaster frigidus* | DGS | 48,325,192 | 9.7 | Tibet, China | PE-Xizang Expedition 2965 (PE) | SRR15691173 | MN577875 |
| *Cotoneaster salicifolius var. henryanus* | DGS | 33,722,386 | 6.7 | Hubei, China | B.B.Liu 2241 (PE) | SRR15691172 | MN577863 |
| *Crataegus laevigata* | WGS | 67,361,324 | 13.5 | Denmark | - | SRR12518628 | OM232780 |
| *Crataegus mollis* | WGS | 283,044,360 | 56.6 | Ontario, Canada | - | SRR3130998 | OM232779 |
| *Crataegus pinnatifida* | RNA-Seq | 52,770,968 | 10.6 | Liaoning, China | - | SRR4048567 | MN102356 |
| *Crataegus rhipidophylla* | WGS | 175,178,342 | 35.0 | Denmark | - | SRR12518826 | OM232778 |
| *Cydonia oblonga* | WGS | 205,274,514 | 41.1 | - | - | SRR3166923 | OM232772 |
| *Dichotomanthes tristaniicarpa* | DGS | 45,866,100 | 9.2 | Yunnan, China | B.B.Liu & F.Zhao 3958 (PE) | SRR15691171 | MN577869 |
| *Docynia delavayi* | RNA-Seq | 94,304,864 | 18.9 | Yunnan, China | XYZ080 (FUS) | SRR15691170 | MN216025 |
| *Eriobotrya japonica* | WGS | 311,656,626 | 62.3 | China | - | SRR10377315 | MN577877 |
| *Eriobotrya seguinii* | DGS | 56,412,898 | 11.3 | Guizhou, China | Z.S.Zhang & Y.T.Zhang 4328 (PE) | SRR15691169 | MN577884 |
| *Eriolobus trilobatus* | RNA-Seq | 143,189,250 | 28.6 | Arnold, USA | 127-2009*A (FUS) | SRR15691168 | KX499858 |
| *Gillenia stipulata* | WGS | 121,927,200 | 24.4 | - | - | SRR3166477 | OM232773 |
| *Kageneckia oblonga* | WGS | 200,549,100 | 40.1 | Santiago, Chile | - | SRR3157483 | OM232774 |
| *Malacomeles denticulata* | RNA-Seq | 28,918,506 | 5.8 | Berkeley, USA | WYQ-14A (FUS) | SRR15691197 | MN068267 |
| *Malus baccata* | WGS | 184,814,116 | 37.0 | Shanxi, China | - | SRR7248844 | OM232791 |
| *Malus florentina* | WGS | 63,939,898 | 12.8 | UK | - | SRR3571157 | OM232784 |
| *Malus fusca* | WGS | 281,596,698 | 56.3 | - | - | SRR1658492 | OM232797 |
| *Malus ioensis* | WGS | 92,592,974 | 18.5 | Illinois, USA | - | SRR3141653 | OM232799 |
| ***Photinia beckii*** | **DGS†** | **54,104,550** | **10.9** | **Yunnan, China** | **L.Y.Wang & H.Z.Feng 1625 (SYS)** | **SRR20278372** | **OP021702*** |
| ***Photinia chihsiniana*** | **DGS†** | **70,608,580** | **14.3** | **Guangxi, China** | **L.Y.Wang & K.W.Xu 1540 (SYS)** | **SRR20278371** | **OP021703*** |
| *Photinia crassifolia* | DGS | 72,077,600 | 14.4 | Guizhou, China | L.Y.Wang & H.Z.Feng 1640 (SYS) | SRR15691190 | MZ984217 |

| | | | | | | | |
|---|---|---|---|---|---|---|---|
| *Photinia glabra* | DGS | 73,353,976 | 14.7 | Jiangxi, China | B.B.Liu P1936-3 (PE) | SRR15691189 | MZ984218 |
| ***Photinia glomerata*** | **DGS†** | **72,736,460** | **14.7** | **Yunnan, China** | **L.Y.Wang 1666 (SYS)** | **SRR20278386** | **OP021704*** |
| ***Photinia integrifolia*** | **DGS†** | **43,624,632** | **8.8** | **Yunnan, China** | **L.Y.Wang et al. 1598 (SYS)** | **SRR20278385** | **OP021705*** |
| ***Photinia kwangsiensis*** | **DGS†** | **55,066,376** | **11.1** | **Guangxi, China** | **L.Y.Wang et al. 1719 (SYS)** | **SRR20278384** | **OP021706*** |
| ***Photinia lanuginosa*** | **DGS†** | **37,230,622** | **7.5** | **Hunan, China** | **Z.C.Luo 730 (PE)** | **SRR20336850** | **MN577890*** |
| ***Photinia lasiogyna 1*** | **DGS†** | **55,230,204** | **11.2** | **Sichuan, China** | **Qinghai-Tibet Expedition 11412 (PE)** | **SRR20278377** | **OP021696*** |
| ***Photinia lasiogyna 2*** | **DGS†** | **50,930,006** | **10.3** | **Sichuan, China** | **Qinghai-Tibet Expedition 13564 (PE)** | **SRR20278376** | **OP021697*** |
| ***Photinia lasiogyna 3*** | **DGS†** | **20,147,234** | **4.1** | **Yunnan, China** | **R.C.Qin 24332 (PE)** | **SRR20278375** | **OP021699*** |
| ***Photinia lasiogyna 4*** | **DGS†** | **20,852,368** | **4.2** | **Yunnan, China** | **780Gongcheng 873 (PE)** | **SRR20336851** | **MK920280*** |
| ***Photinia lochengensis*** | **DGS†** | **29,682,686** | **6.0** | **Guangxi, China** | **G.R.Long 89009 (PE)** | **SRR20336849** | **MN577888*** |
| ***Photinia loriformis*** | **DGS†** | **64,458,316** | **13.0** | **Yunnan, China** | **L.Y.Wang 1671 (SYS)** | **SRR20278383** | **OP021707*** |
| ***Photinia prionophylla*** | **DGS†** | **38,691,128** | **7.8** | **Yunnan, China** | **Z.D.Fang et al. 20-427 (PE)** | **SRR20336848** | **MN577891*** |
| ***Photinia prunifolia*** | **DGS†** | **72,079,706** | **14.6** | **Zhejiang, China** | **B.B.Liu et al. 3233 (PE)** | **SRR20278382** | **OP021708*** |
| ***Photinia raupingensis*** | **DGS†** | **75,266,998** | **15.2** | **Guangdong, China** | **L.Y.Wang 1788 (SYS)** | **SRR20278381** | **OP021709*** |
| ***Photinia serratifolia*** | **DGS†** | **73,522,912** | **14.9** | **Zhejiang, China** | **B.B.Liu et al. 2887 (PE)** | **SRR20278380** | **OP021710*** |
| ***Photinia stenophylla*** | **DGS†** | **71,763,530** | **14.5** | **Guizhou, China** | **L.Y.Wang 1648 (SYS)** | **SRR20278379** | **OP021711*** |
| *Pourthiaea arguta* | DGS | 99,096,326 | 19.8 | Yunnan, China | E.D.Liu LED6340 (KUN) | SRR15691188 | MT249048 |
| *Pourthiaea pilosicalyx* | DGS | 59,852,032 | 12.0 | Guangxi, China | B.B.Liu 2131 (PE) | SRR15691186 | MT249045 |
| *Pourthiaea zhejiangensis* | DGS | 103,871,714 | 20.8 | Zhejiang, China | L.Y.Wang T29509 (SYS) | SRR15691185 | MZ984216 |
| *Pseudocydonia sinensis* | DGS | 36,327,950 | 7.3 | Beijing, China | B.B.Liu 3981 (PE) | SRR15691184 | MN577871 |
| *Pyracantha coccinea* | WGS | 100,901,170 | 20.2 | Bethesda, USA | IRGN X329LVHRXG | SRR13004386 | OM232776 |
| *Pyrus communis* | WGS | 247,493,422 | 49.5 | Angers, France | - | SRR10030308 | MN577870 |
| *Pyrus pyrifolia* | RNA-Seq | 112,490,888 | 22.5 | Sanming, China | - | SRR10415525 | AP012207 |
| *Rhaphiolepis ferruginea* | DGS | 47,935,428 | 9.6 | Guangxi, China | B.B.Liu P1900-3 (PE) | SRR15691183 | MN577866 |
| *Rhaphiolepis umbellata* | DGS | 47,845,048 | 9.6 | Zhejiang, China | B.B.Liu 1951 (PE) | SRR15691182 | MN577868 |
| *Sorbus americana* | DGS | 39,540,022 | 7.9 | Tennessee, USA | Philippe et al. 42803 (PE) | SRR15691181 | MZ984219 |
| *Sorbus commixta* | RNA-Seq | 25,003,474 | 5.0 | Hubei, China | XYZ039 (FUS) | SRR15691180 | MK920288 |
| *Sorbus ferruginea* | DGS | 41,054,942 | 8.2 | Yunnan, China | Hongheshui Expedition 2204 (PE) | SRR15691179 | MZ984220 |
| *Sorbus hemsleyi* | DGS | 47,864,196 | 9.6 | Sichuan, China | Jeon et al. SI1557 (PE) | SRR15691178 | MZ984221 |
| *Sorbus matsumurana* | DGS | 40,408,954 | 8.1 | Tottori, Japan | Inoue 100634 (PE) | SRR15691177 | MZ984222 |

| | | | | | | | |
|---|---|---|---|---|---|---|---|
| *Sorbus pteridophylla* | DGS | 79,221,176 | 15.8 | Tibet, China | PE-Xizang Expedition 3354 (PE) | SRR15691175 | MZ984223 |
| *Sorbus slavnicensis* | DGS | 34,381,772 | 6.9 | Croatia | V.Mikolas et al. s.n. (PE) | SRR15691174 | MZ984224 |
| ***Stranvaesia bodinieri 1*** | **DGS†** | **74,500,164** | **15.1** | **Jiangxi, China** | **B.B.Liu P1941-3 (PE)** | **SRR20278388** | **OP021698*** |
| ***Stranvaesia bodinieri 2*** | **DGS†** | **38,270,610** | **7.7** | **Guangxi, China** | **Y.M.Wang 322 (PE)** | **SRR20278387** | **OP021695*** |
| ***Stranvaesia nussia*** | **DGS†** | **50,159,118** | **10.1** | **USA** | **D.H.Nicolson 2719 (US)** | **SRR20336852** | **MK920284*** |
| ***Stranvaesia oblanceolata 1*** | **DGS†** | **34,364,088** | **6.9** | **Yunnan, China** | **H.Wang 3666 (PE)** | **SRR20278378** | **OP021694*** |
| ***Stranvaesia oblanceolata 2*** | **DGS†** | **58,847,430** | **11.9** | **Yunnan, China** | **G.Forrest 11855 (PE)** | **SRR20278374** | **OP021700*** |
| ***Stranvaesia oblanceolata 3*** | **DGS†** | **58,788,222** | **11.9** | **Yunnan, China** | **P.Y.Mao 06729 (PE)** | **SRR20278373** | **OP021701*** |
| *Vauquelinia californica* | WGS | 195,790,058 | 39.2 | - | - | SRR3130625 | OM232777 |

† indicates that the sample was sequenced for this study. DGS, deep genome skimming; WGS; whole-genome resequencing; RNA-Seq: transcriptome;

* indicates that the plastome was assembled for this study.

FUS: Herbarium of Fudan University

PE: Chinese National Herbarium

SYS: Herbarium of Sun Yat-sen University

KUN: Kunming Institute of Botany, Chinese Academy of Sciences

US: Smithsonian Institution
