## Supplementary_material for "Nightmare or delight: taxonomic circumscription meets reticulate evolution in the phylogenomic era": SUPPLEMENTARY.TABLE.S2.pdf

Table S2. The number of pre-filtered and post-filtered sequences for each sample in this study

| Names | Pre-filtered sequences | Post-filtered sequences |
|---|---|---|
| *Amelanchier alnifolia* | 684 | 332 |
| *Aronia melanocarpa* | 658 | 325 |
| *Chaenomeles japonica* | 756 | 379 |
| *Chaenomeles speciosa* | 792 | 396 |
| *Cormus domestica* | 724 | 357 |
| *Cotoneaster frigidus* | 780 | 406 |
| *Cotoneaster salicifolius var. henryanus* | 687 | 335 |
| *Crataegus laevigata* | 797 | 395 |
| *Crataegus mollis* | 801 | 404 |
| *Crataegus pinnatifida* | 728 | 356 |
| *Crataegus rhipidophylla* | 801 | 405 |
| *Cydonia oblonga* | 800 | 400 |
| *Dichotomanthes tristaniicarpa* | 742 | 364 |
| *Docynia delavayi* | 763 | 384 |
| *Eriobotrya japonica* | 800 | 403 |
| *Eriobotrya seguinii* | 800 | 403 |
| *Eriolobus trilobatus* | 718 | 350 |
| *Gillenia stipulata* | 800 | 405 |
| *Kageneckia oblonga* | 800 | 404 |
| *Malacomeles denticulata* | 706 | 348 |
| *Malus baccata* | 800 | 407 |
| *Malus florentina* | 801 | 408 |
| *Malus fusca* | 801 | 412 |
| *Malus ioensis* | 799 | 410 |
| *Photinia beckii* | 797 | 401 |
| *Photinia chihsiniana* | 798 | 410 |
| *Photinia crassifolia* | 798 | 410 |
| *Photinia glabra* | 796 | 404 |
| *Photinia glomerata* | 799 | 417 |
| *Photinia integrifolia* | 704 | 339 |
| *Photinia kwangsiensis* | 797 | 398 |
| *Photinia lanuginosa* | 735 | 355 |
| *Photinia lasiogyna 1* | 791 | 392 |
| *Photinia lasiogyna 2* | 793 | 391 |
| *Photinia lasiogyna 3* | 304 | - |
| *Photinia lasiogyna 4* | 527 | - |
| *Photinia lochengensis* | 636 | 294 |
| *Photinia loriformis* | 798 | 414 |
| *Photinia prionophylla* | 647 | 307 |
| *Photinia prunifolia* | 767 | 378 |
| *Photinia raupingensis* | 794 | 398 |
| *Photinia serratifolia* | 767 | 393 |
| *Photinia stenophylla* | 796 | 403 |

| | | |
|---|---|---|
| *Pourthiaea arguta* | 801 | 401 |
| *Pourthiaea pilosicalyx* | 779 | 388 |
| *Pourthiaea zhejiangensis* | 799 | 401 |
| *Pseudocydonia sinensis* | 631 | 294 |
| *Pyracantha coccinea* | 801 | 403 |
| *Pyrus communis* | 801 | 405 |
| *Pyrus pyrifolia* | 750 | 376 |
| *Rhaphiolepis ferruginea* | 714 | 352 |
| *Rhaphiolepis umbellata* | 726 | 360 |
| *Sorbus americana* | 743 | 366 |
| *Sorbus commixta* | 712 | 365 |
| *Sorbus ferruginea* | 775 | 391 |
| *Sorbus hemsleyi* | 784 | 393 |
| *Sorbus matsumurana* | 758 | 390 |
| *Sorbus pteridophylla* | 528 | 245 |
| *Sorbus slavnicensis* | 657 | 314 |
| *Stranvaesia bodinieri 1* | 796 | 408 |
| *Stranvaesia bodinieri 2* | 737 | 354 |
| *Stranvaesia nussia* | 523 | 232 |
| *Stranvaesia oblanceolata 1* | 620 | 277 |
| *Stranvaesia oblanceolata 2* | 799 | 401 |
| *Stranvaesia oblanceolata 3* | 800 | 402 |
| *Vauquelinia californica* | 801 | 399 |
