## Supplementary figures and images for "Nightmare or delight: taxonomic circumscription meets reticulate evolution in the phylogenomic era"

### SUPPLEMENTARY.FIG.S1.PDF.pdf

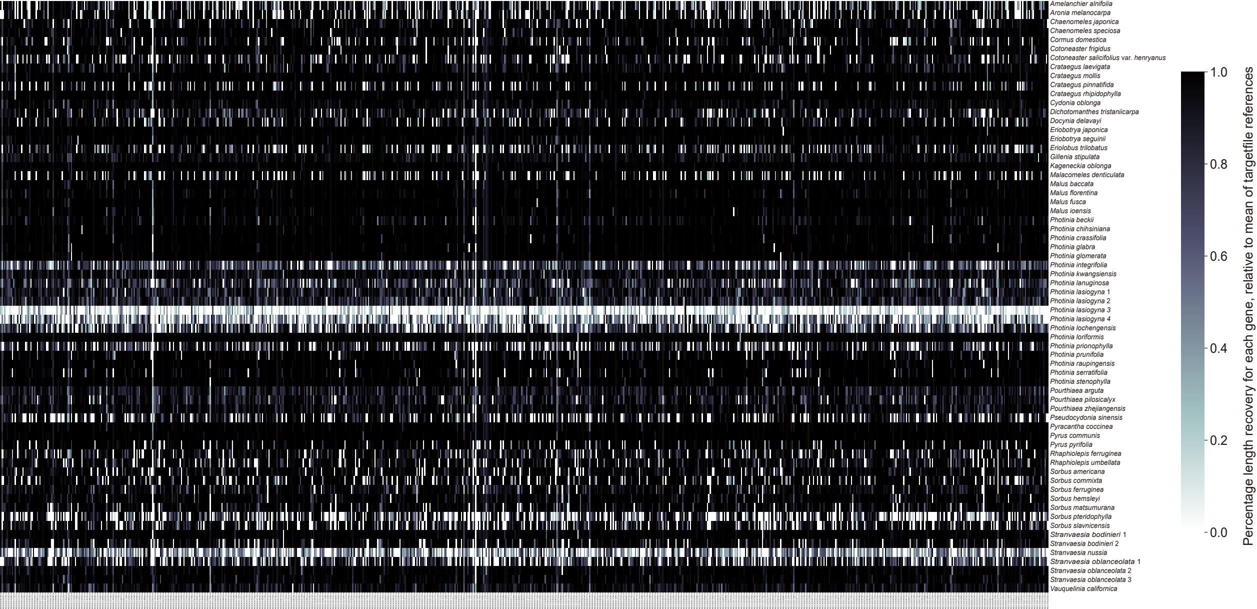

### SUPPLEMENTARY.FIG.S2.PDF.pdf

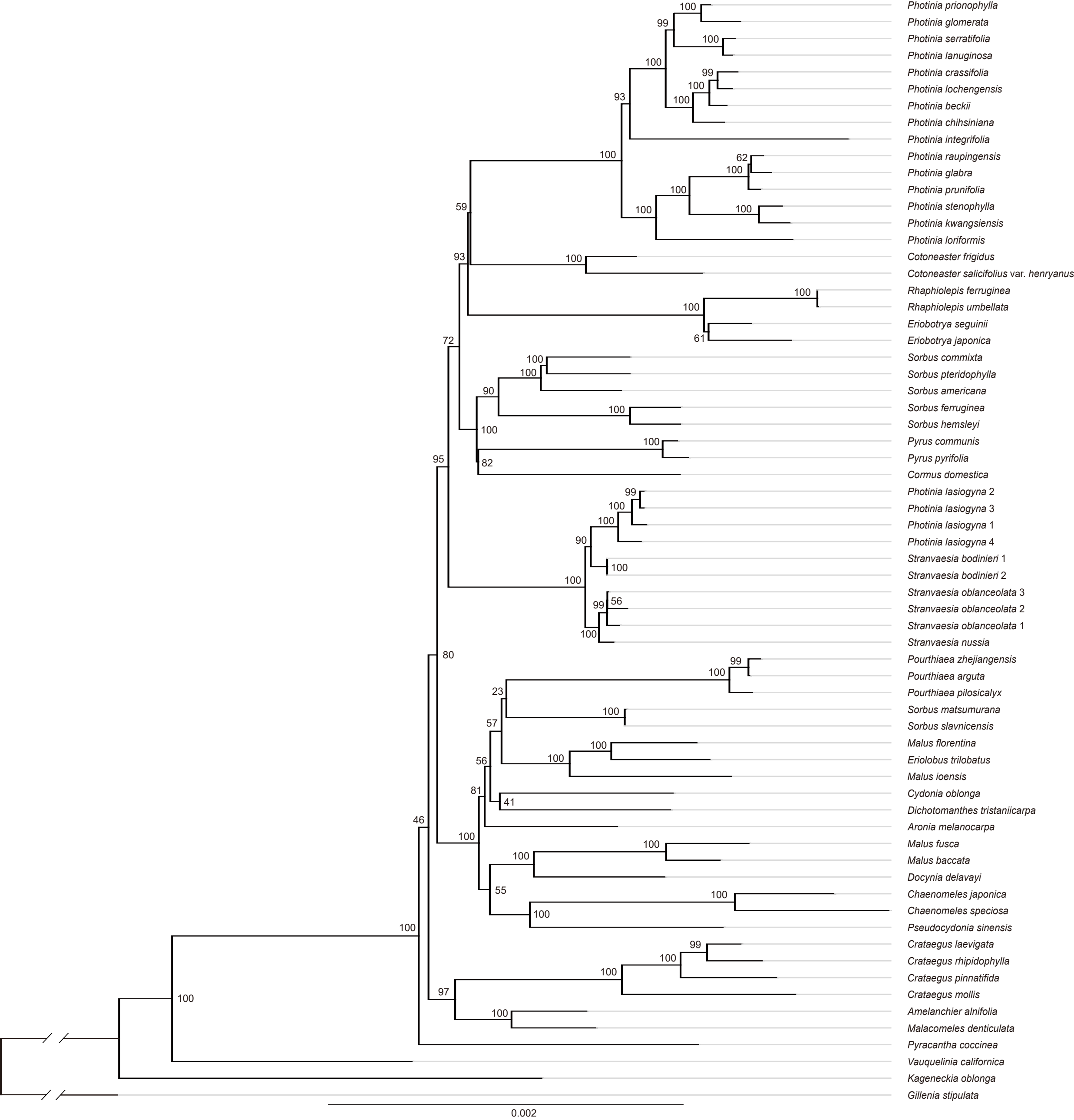

### SUPPLEMENTARY.FIG.S3.PDF.pdf

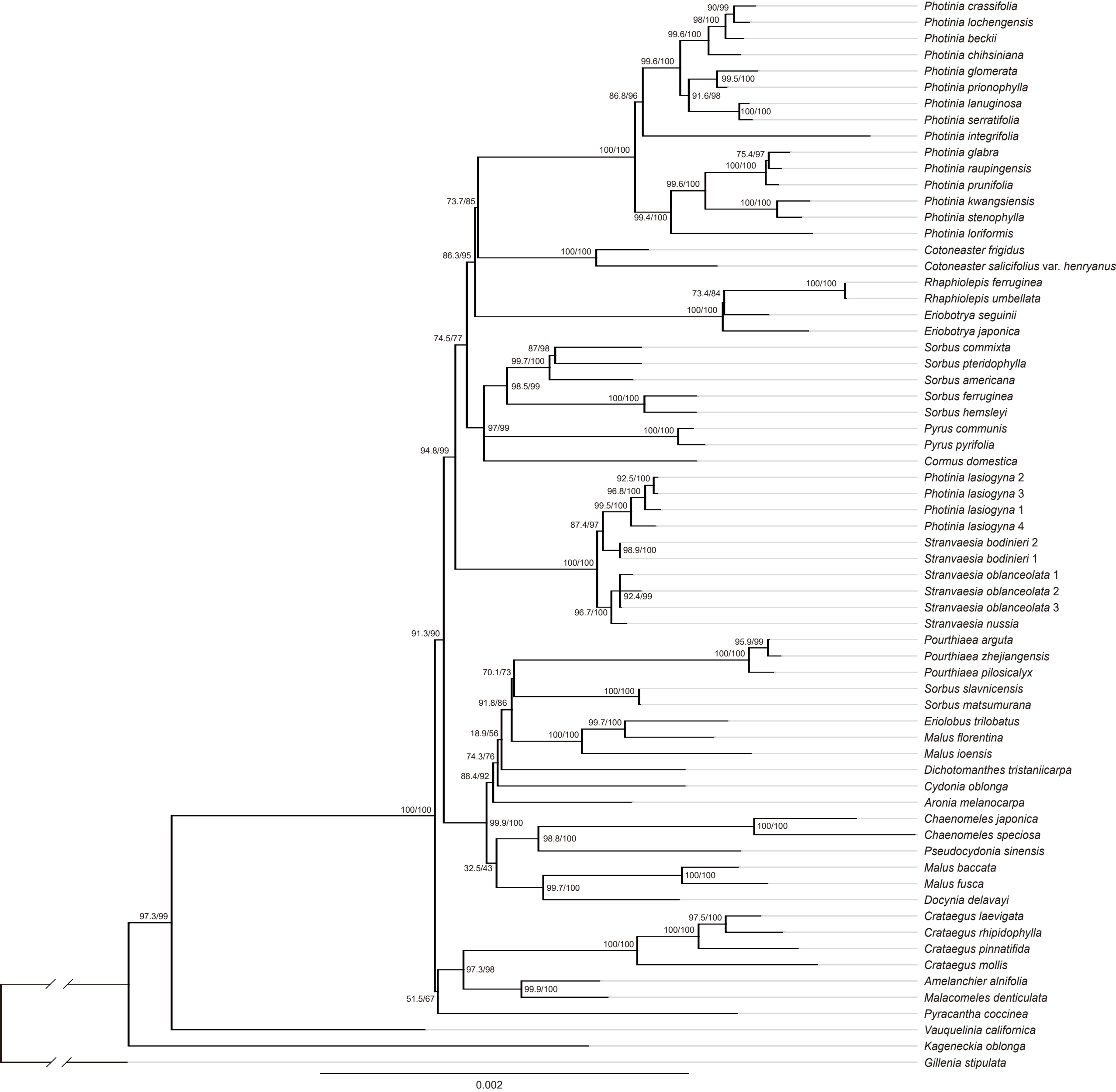

### SUPPLEMENTARY.FIG.S4.PDF.pdf

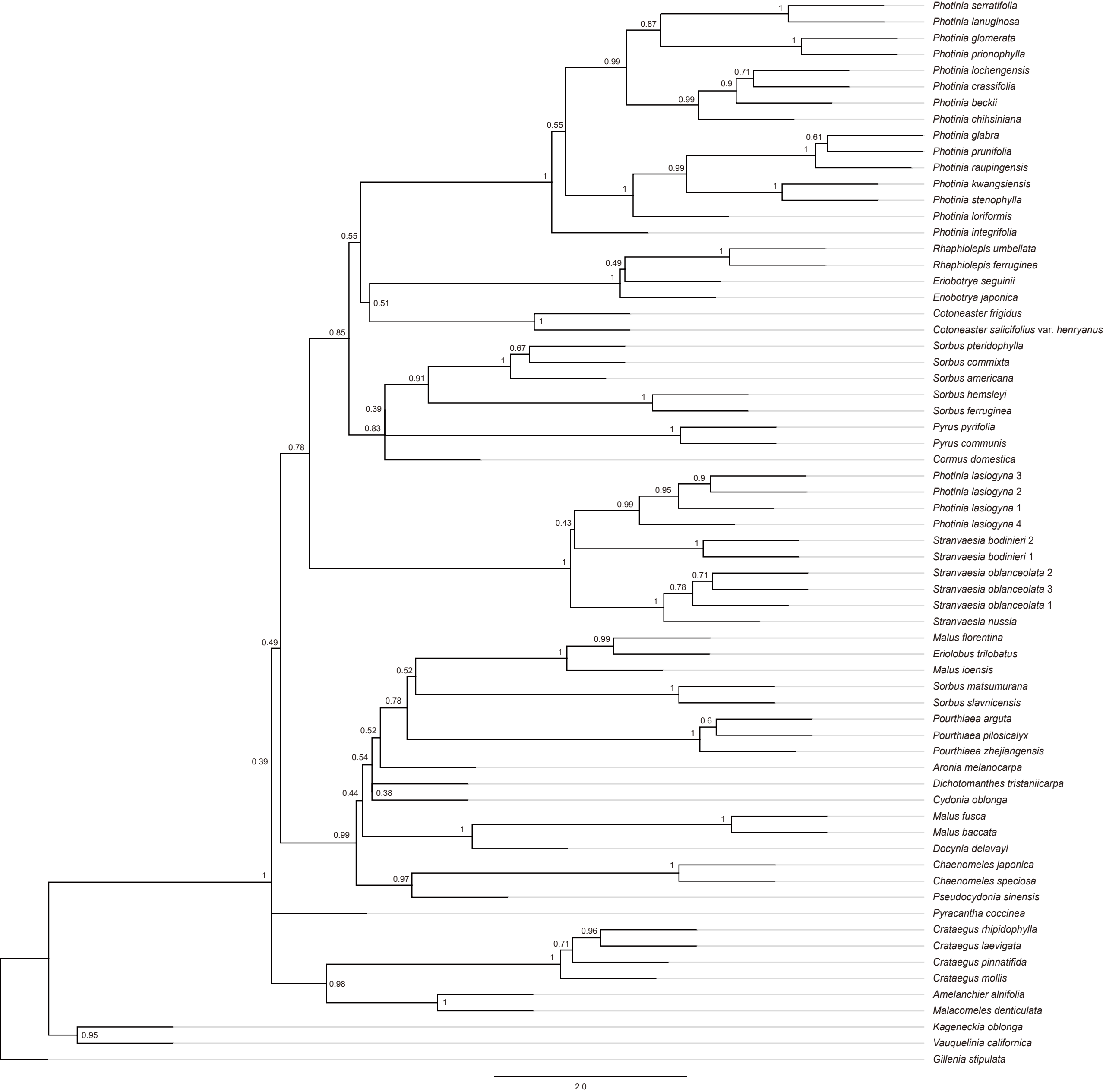

### SUPPLEMENTARY.FIG.S5.PDF.pdf

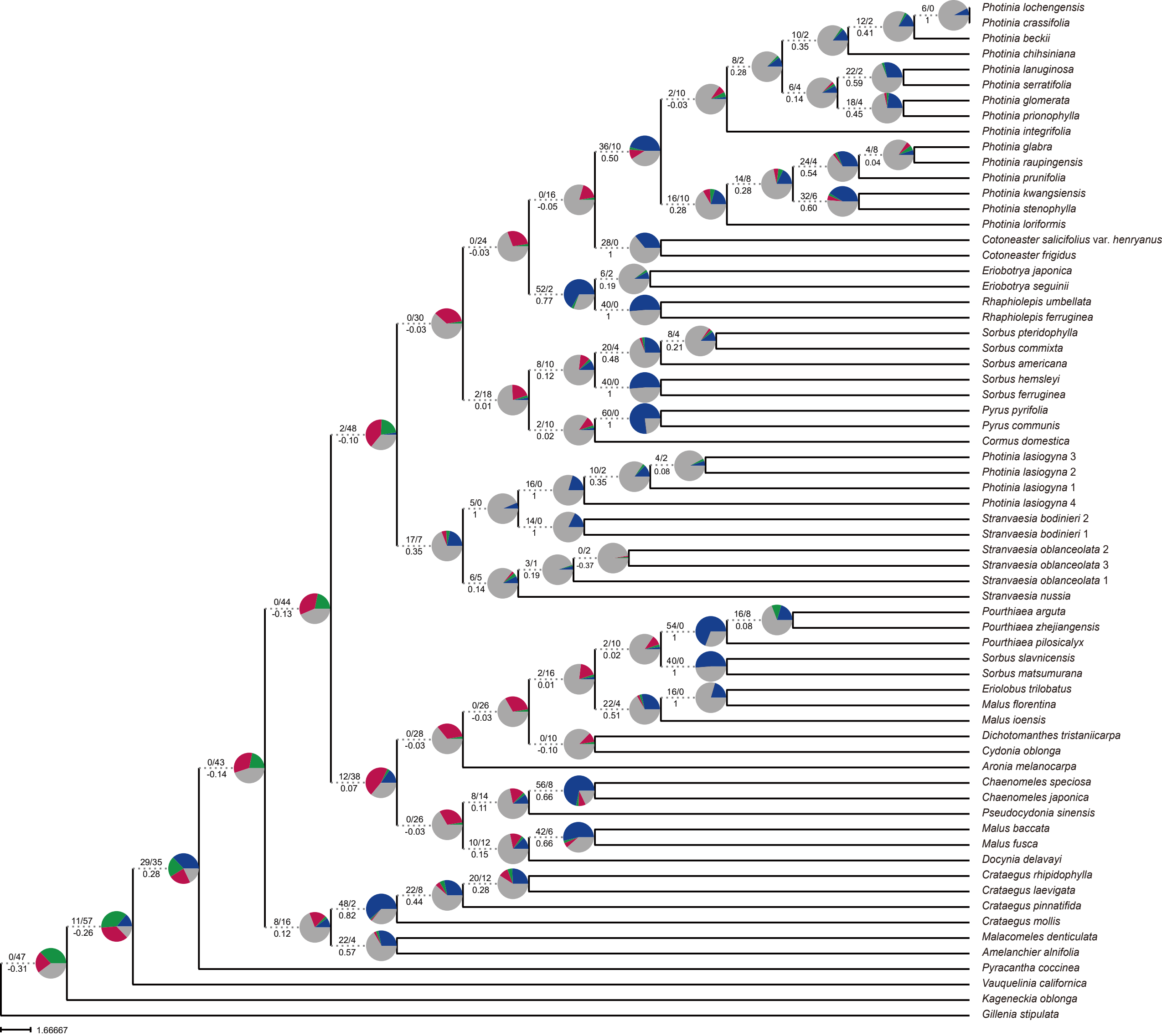

### SUPPLEMENTARY.FIG.S6.PDF.pdf

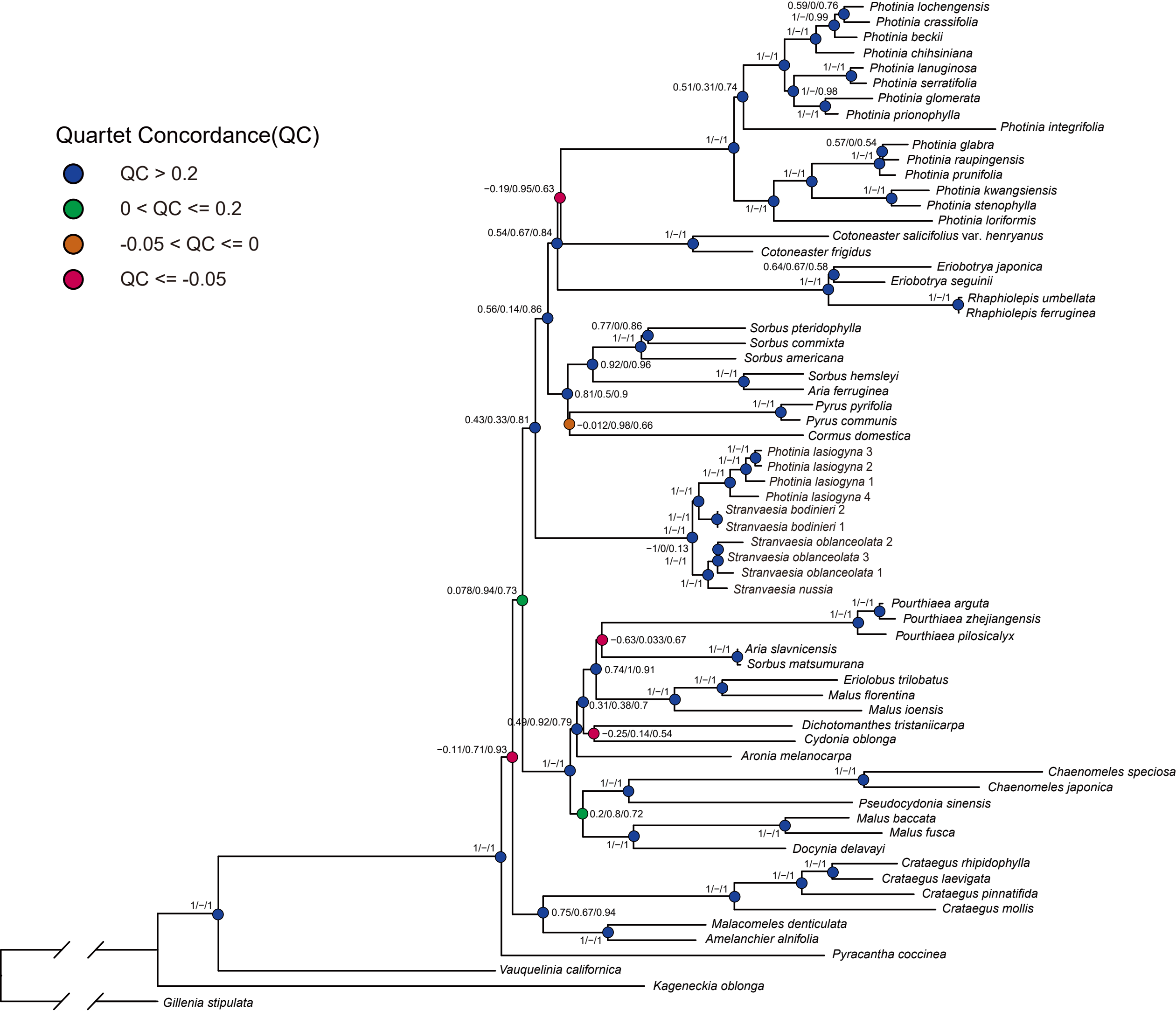

### SUPPLEMENTARY.FIG.S7.PDF.pdf

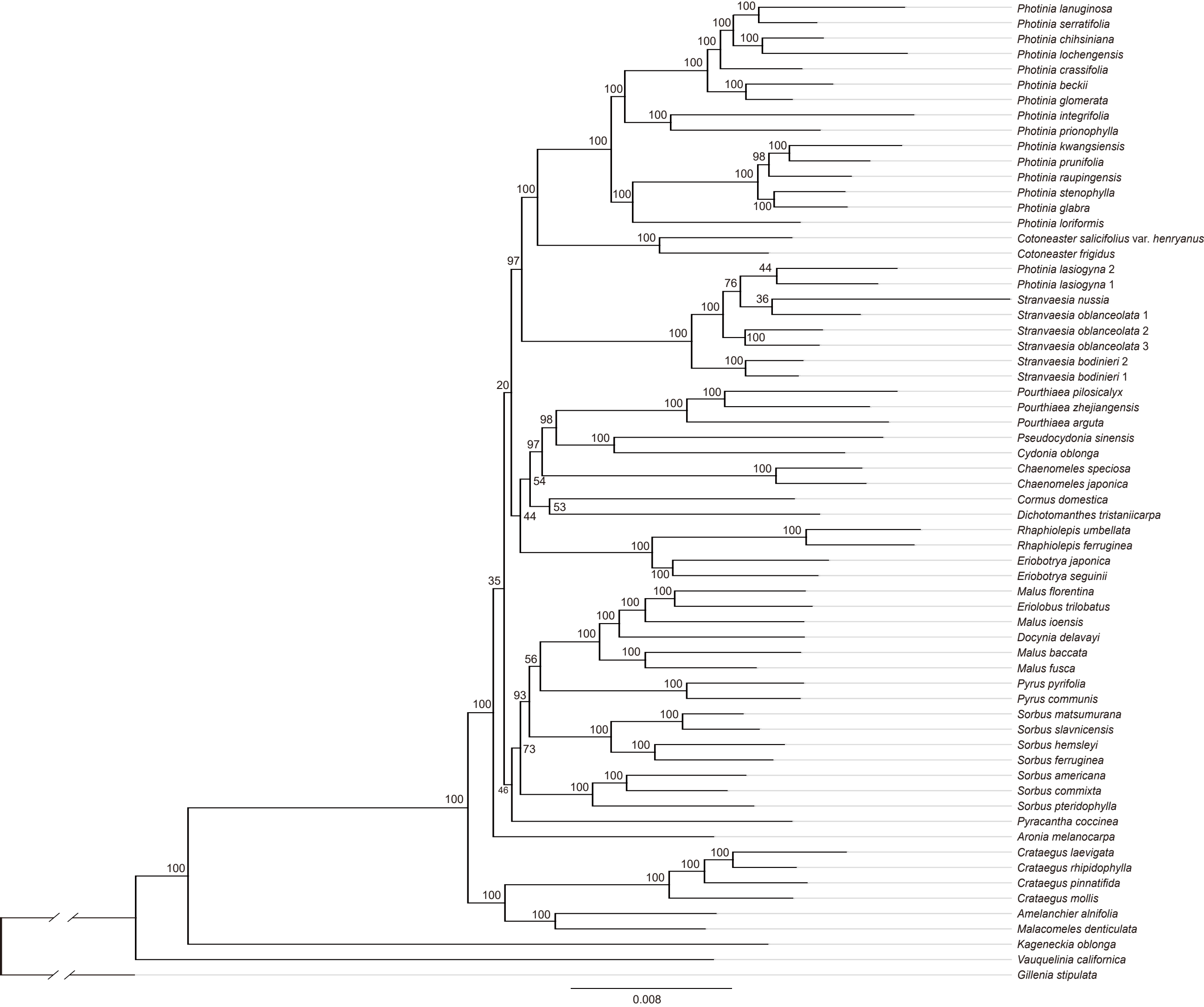

### SUPPLEMENTARY.FIG.S8.PDF.pdf

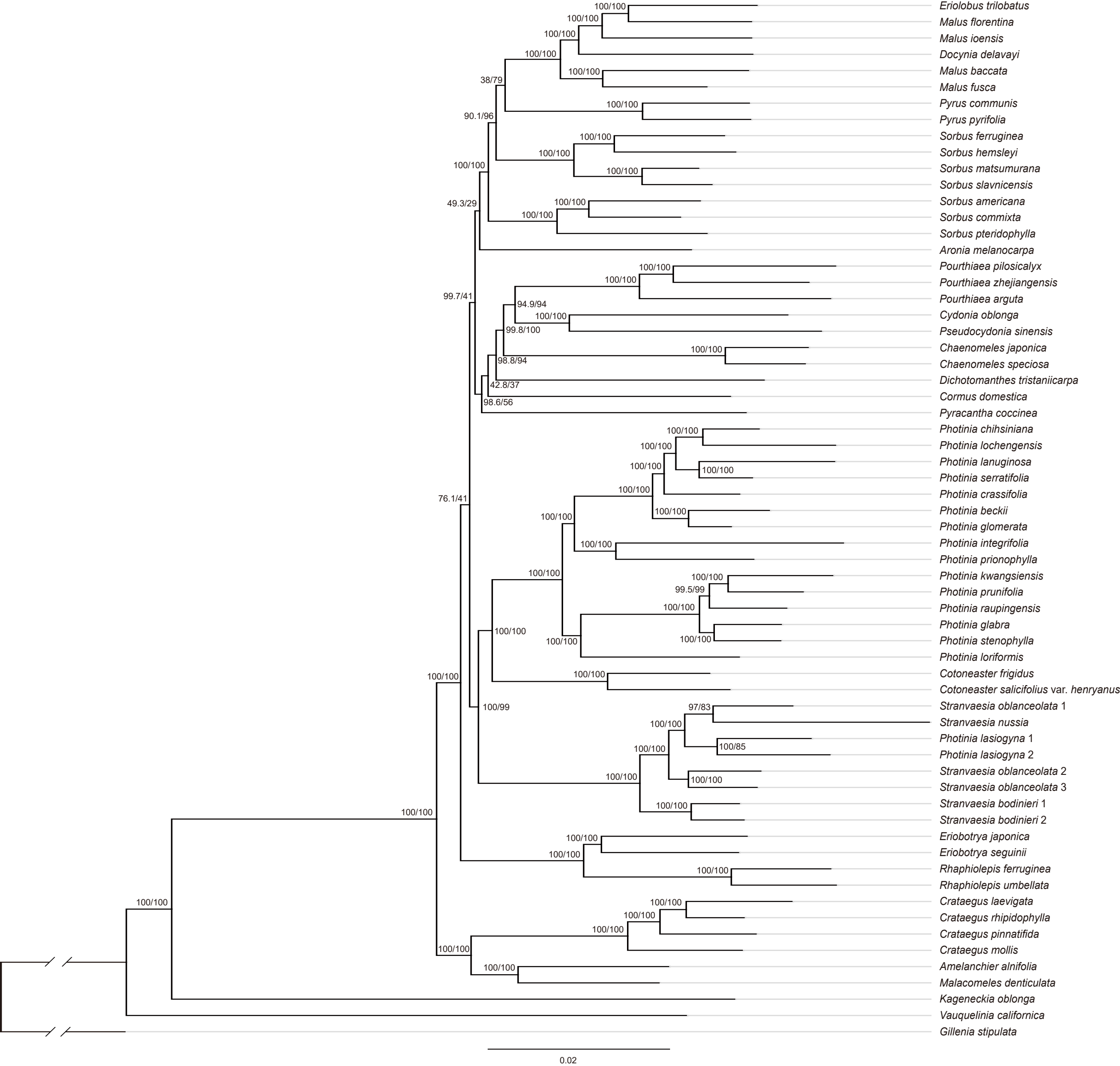

### SUPPLEMENTARY.FIG.S9.PDF.pdf

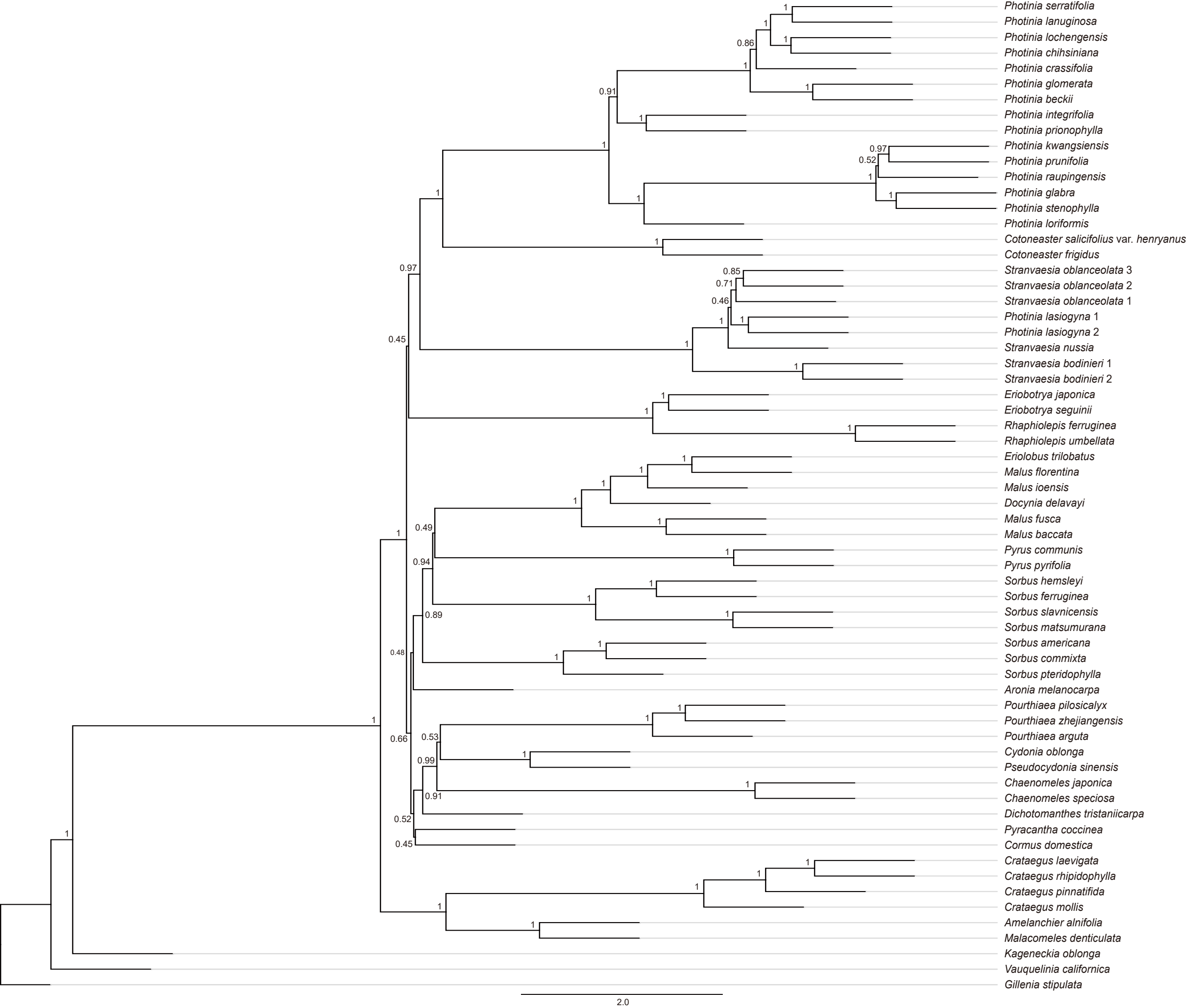

### SUPPLEMENTARY.FIG.S10.PDF.pdf

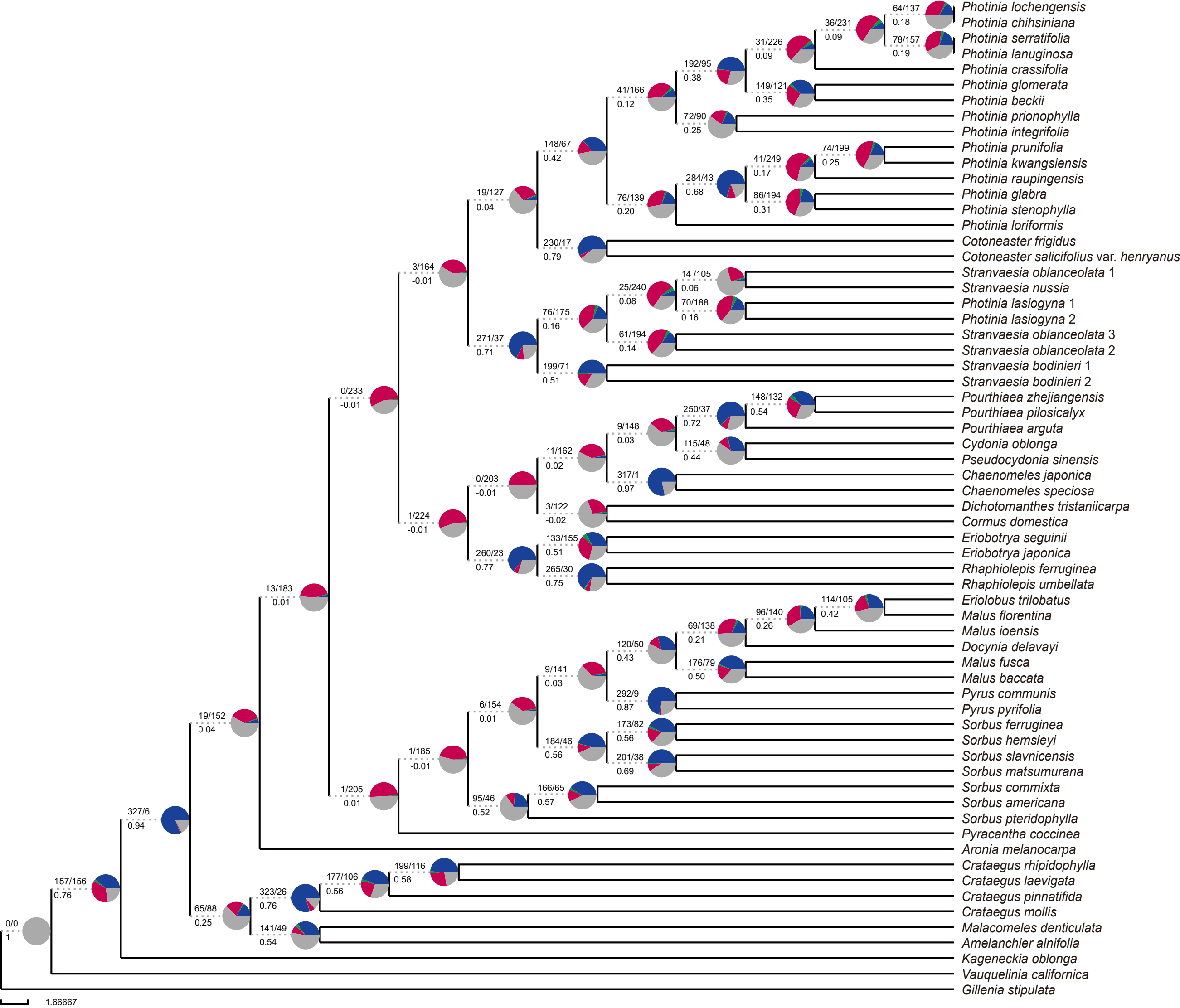

### SUPPLEMENTARY.FIG.S11.PDF.pdf

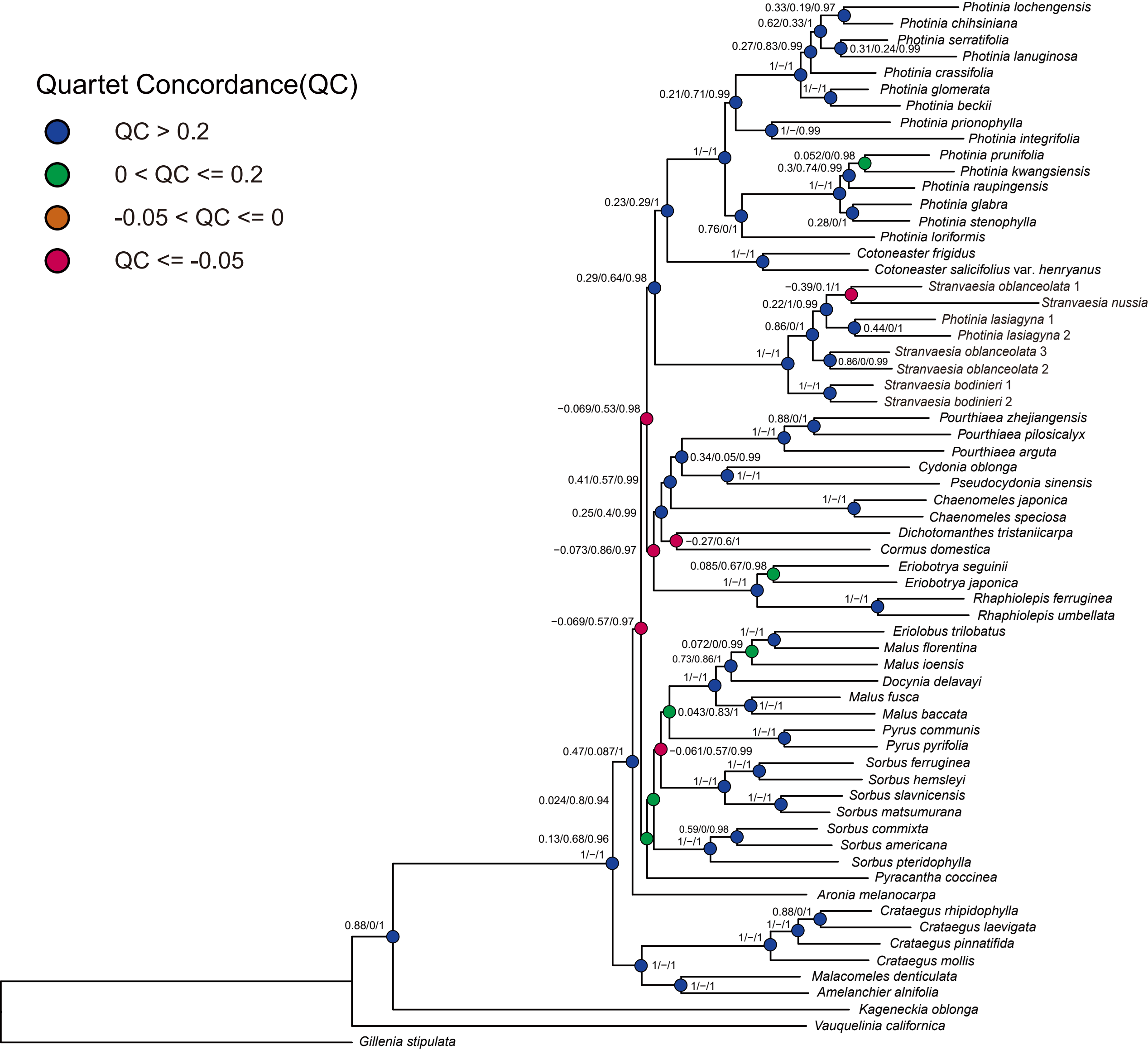

### SUPPLEMENTARY.FIG.S12.PDF.pdf

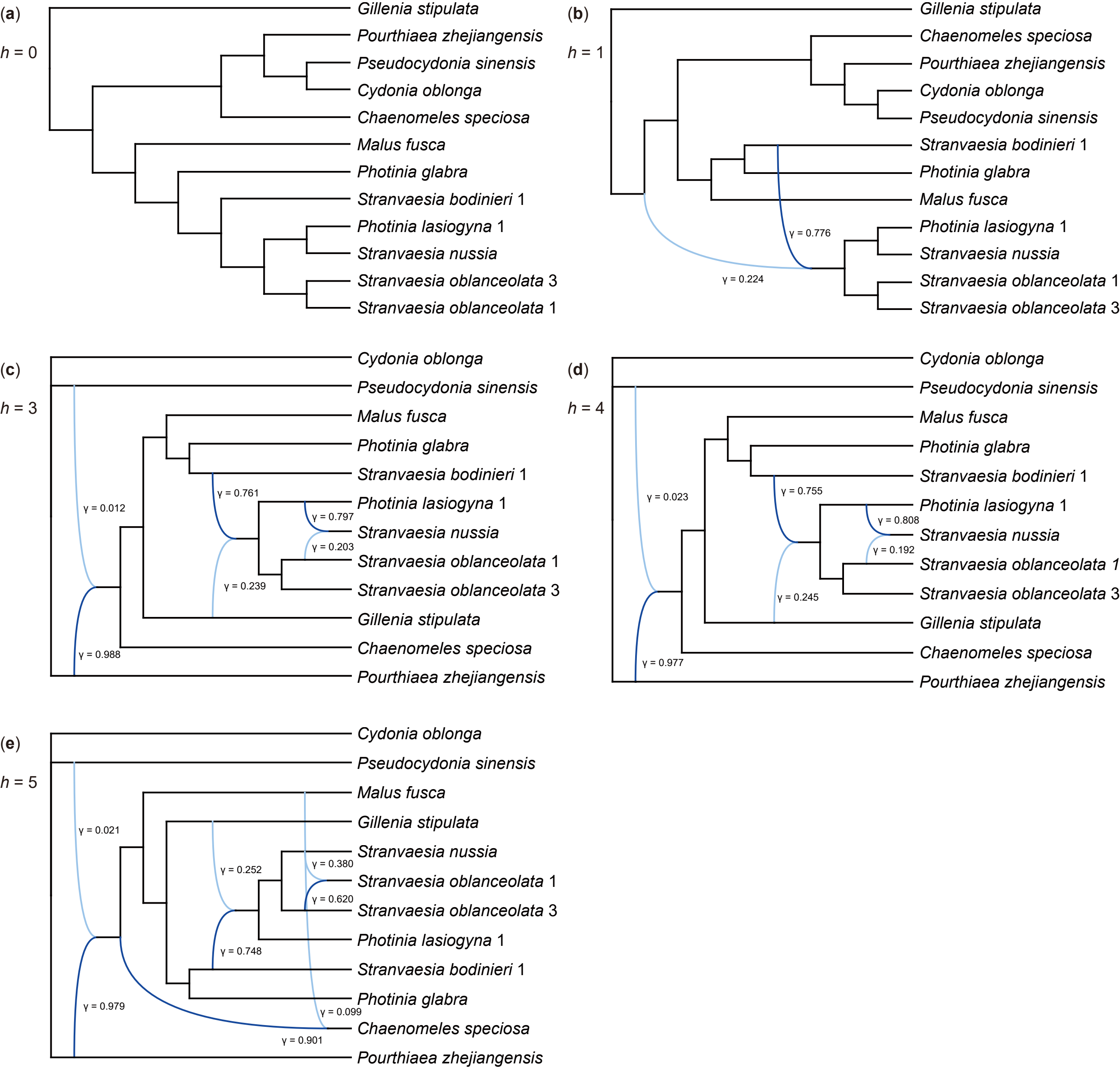

### SUPPLEMENTARY.FIG.S13.PDF.pdf

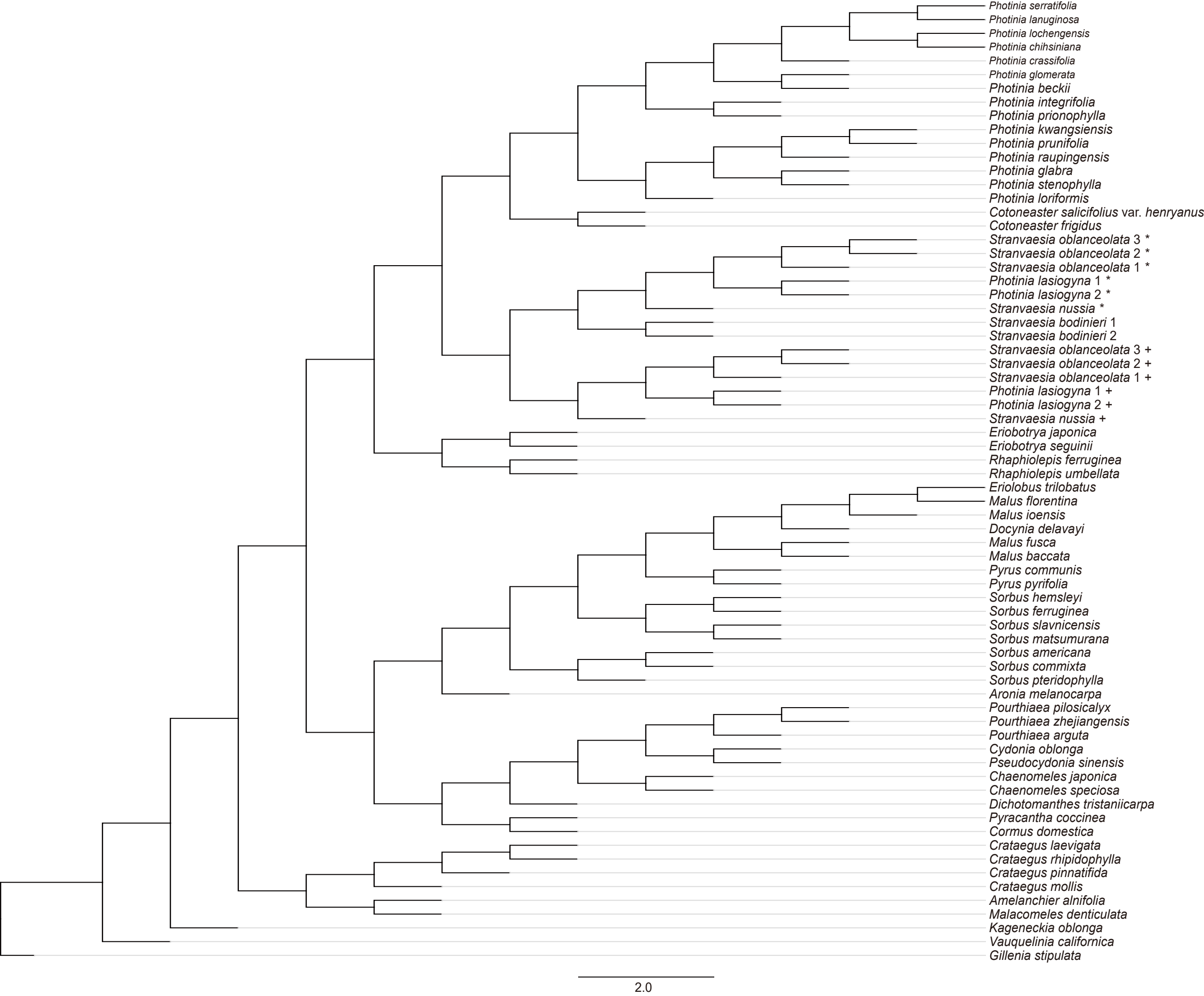

### SUPPLEMENTARY.FIG.S14.PDF.pdf

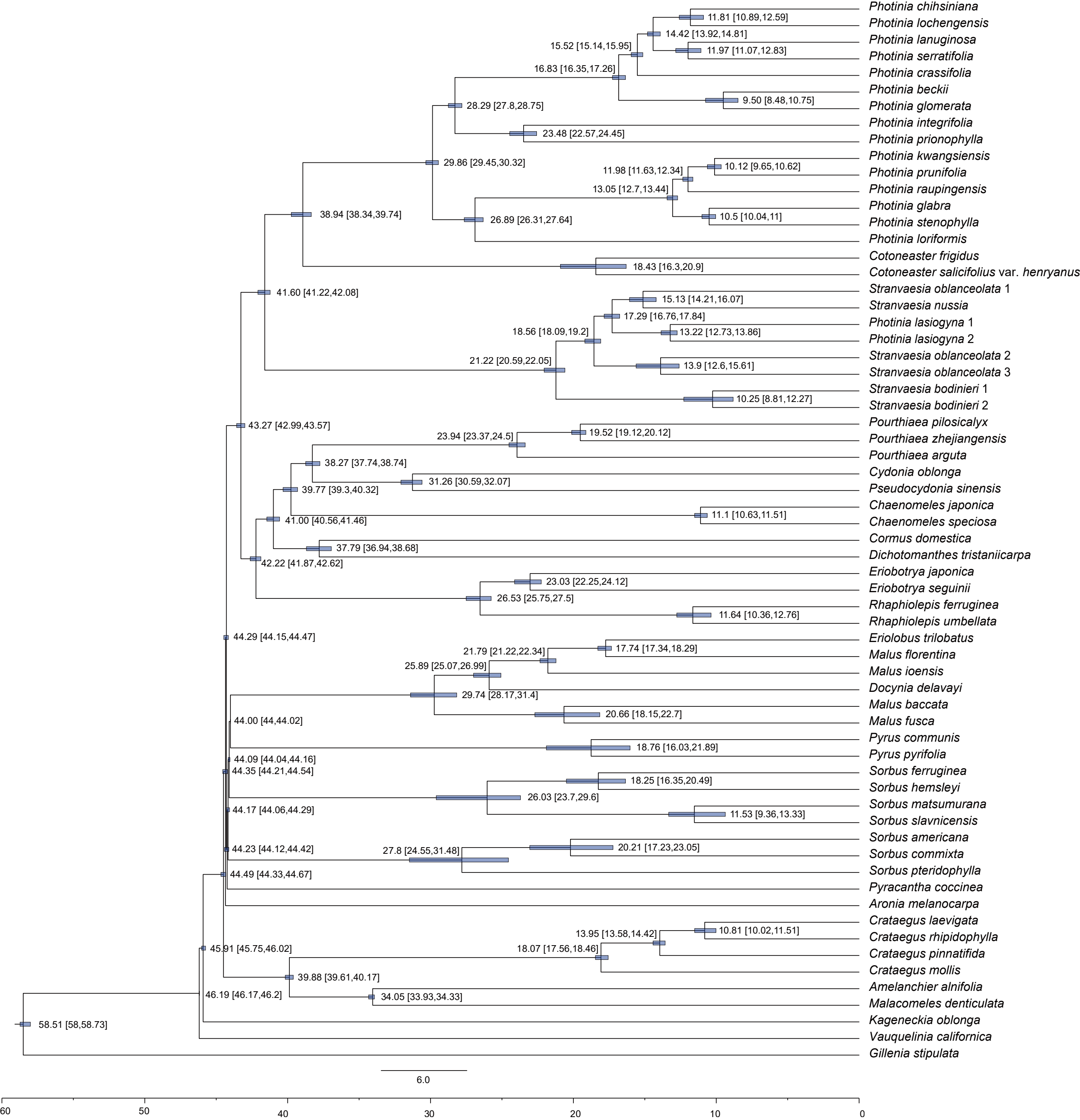

### SUPPLEMENTARY.FIG.S15.PDF.pdf

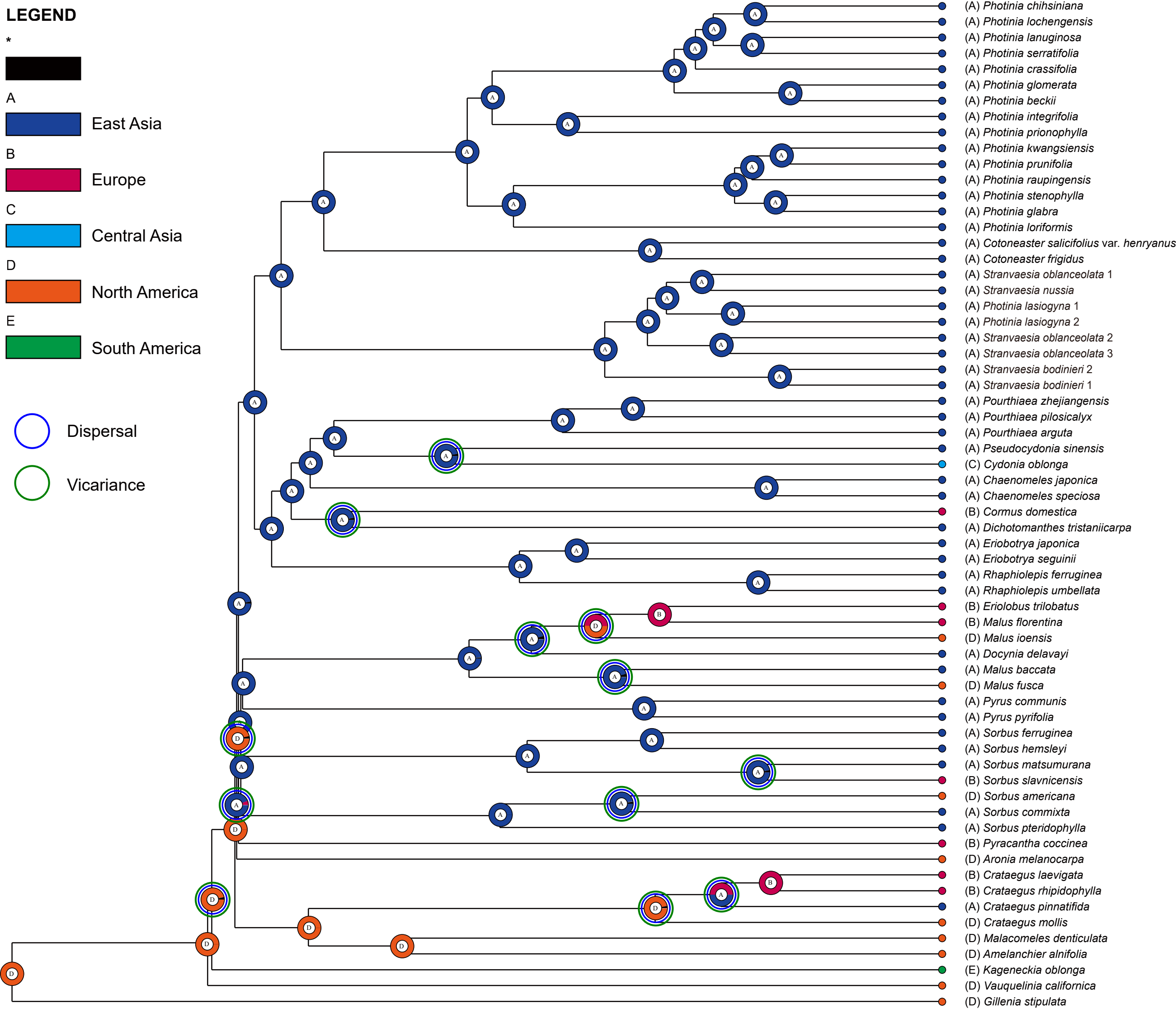
